# Respiration-Deficient Cells Require Pyruvate Carboxylase to Suppress Asparagine Auxotrophy

**DOI:** 10.64898/2026.08.12.744280

**Authors:** Ruobing Cui, Keun Woo Ryu, Yi Fu, Ziad Bakouny, Dayi Li, Tamar Kavlashvili, Agnel Sfeir, Craig B. Thompson

## Abstract

Mutations in mitochondrial DNA (mtDNA) compromise ETC activity and impair oxidative phosphorylation. Since eukaryotic cells contain multiple copies of mtDNA, the resulting phenotype depends on the proportion of mutant mitochondrial genomes (the heteroplasmy level). Using isogenic cell lines carrying similar mtDNA deletions, a linear decline in cellular respiration was observed as mitochondrial DNA heteroplasmy increased. Despite this, cellular redox imbalance did not change until heteroplasmy exceeded 50%. As heteroplasmy increased past 70%, cells also exhibited an integrated stress response (ISR) and impaired translation was observed. These defects were reversed by either addition of asparagine or overexpression of pyruvate carboxylase (PC). The dependence on exogenous asparagine in other respiration-deficient cells was found to correlate inversely with the PC expression level. For example, patient-derived thyroid tumor cells, harboring high heteroplasmy for a Complex I mtDNA mutation and low levels of PC, exhibited asparagine auxotrophy, and L-asparaginase treatment suppressed tumor growth. Together, these findings demonstrate a role for mitochondrial pyruvate carboxylase in cellular asparagine synthesis under conditions of compromised respiratory activity.

## Main

Mitochondria are central hubs of cellular metabolism, supporting both bioenergetic and biosynthetic processes^1,2^. Mitochondrial oxidation of bioenergetic substrates supports electron transport chain activity and ATP generation through oxidative phosphorylation (OXPHOS). Electron transport chain (ETC) activity also supports the synthesis of pyrimidines by assimilating the electrons generated during the conversion of dihydroorotate to orotate by the mitochondrial inner membrane protein, dihydroorotate dehydrogenase (DHODH)^3,4^. In addition to these ETC-dependent syntheses, mitochondria also play critical roles in iron-sulfur complex assembly, heme biosynthesis, and the production of lipids and nonessential amino acids^2,5–9^.

Evolutionary evidence suggests mitochondria originated from α-proteobacteria and evolved into endosymbiotic organelles^10,11^. Most eukaryotic cells still harbor mitochondria that retain circular double-stranded DNA (dsDNA). In humans, mtDNA encodes 13 proteins and 24 RNA components (2 rRNAs and 22 tRNAs), and each is required for the assembly and function of the ETC. Individual mammalian cells contain hundreds to thousands of mtDNA molecules^12,13^. Early studies of the role of mtDNA in mammalian cells centered on 143B Rho^0^ cells which completely lack mtDNA. The 143B Rho^0^ cell line was created by transforming an established human osteosarcoma cell line with the murine oncogene Ki-ras and then subjecting the cells to long-term exposure with ethidium bromide to eliminate mitochondrial DNA^14,15^. First reported in 1989, 143B Rho^0^ cells can be stably passaged in medium supplemented with uridine to support pyrimidine biosynthesis and pyruvate to prevent cytosolic reductive stress and maintain glycolytic ATP production^15^.

Mitochondria have a limited capacity to repair mtDNA and most damaged mtDNA is cleared by exonuclease degradation^16,17^. Nevertheless, mtDNA is susceptible to mutation^12,13^. Over the last several decades, a wide spectrum of diseases linked to mtDNA mutations have been identified, including MELAS (Mitochondrial Encephalopathy, Lactic Acidosis, and Stroke-like episodes), Leigh syndrome, progressive external ophthalmoplegia (PEO), Pearson syndrome, Kearns-Sayre syndrome (KSS), and certain cancers. Pathogenic mtDNA mutations include both point mutations and insertions/deletions^18–21^. When these mutations compromise ETC function, the severity of the illness is determined by both the tissue in which the mutation arises and the ratio of the mutant mtDNA copy number to the total mtDNA copy number in the cell (% heteroplasmy)^22^. Heteroplasmy levels can vary widely across tissues and disease contexts, ranging from near-wildtype to near-homoplasmy. In cancer, a subset of mutations - particularly truncating variants in OXPHOS genes - can clonally expand to high or near-homoplasmic levels. Such high-heteroplasmy mutations are enriched in specific tumor types, especially thyroid, renal, and colorectal cancers, raising the question of whether cancers with high heteroplasmic mtDNA mutations depend on additional exogenous substrates to maintain their ability to proliferate^23–25^, beyond the known requirement for exogenous uridine and pyruvate^15^. For example, studies of 143B cells have demonstrated that exogenous aspartate can suppress the requirement of these cancer cells for exogenous pyruvate^26,27^. 143B Rho^0^ cells have been used to propagate mutant mtDNA from patients with mitochondrial diseases through the creation of cybrids^28^. While this system has been instrumental in characterizing the consequences of mtDNA mutations on mitochondrial and cellular function, neither the contributions of the nuclear DNA mutations present in the 143B host cell nor the role of mtDNA heteroplasmy in the observed phenotypes have been explored. To directly assess the impact of mtDNA heteroplasmy in a non-transformed genetic background, and given that certain cancers harbor high-heteroplasmy mtDNA mutations despite severely compromised cellular respiration, we studied non-transformed human retinal pigment epithelial (RPE) cell lines carrying a 3.5 kb deletion of mtDNA at varying heteroplasmy^29^. This deletion encompasses three mitochondrial tRNA genes, three complex I subunits, two ATP synthase components, and two cytochrome oxidase subunits—recapitulating the common deletion observed in several mitochondrial disease syndromes.

Beginning with these isogenic cell lines, we investigated the relationship between mtDNA heteroplasmy levels and the contribution of mitochondria to cellular ATP production, reductive stress control, and the production of metabolites necessary for maintaining cell survival and proliferation.

## Results

### Mitochondrial DNA heteroplasmic cells exhibit defective oxidative metabolism

To explore how varying degrees of mitochondrial DNA heteroplasmy affect cell viability and proliferation, a set of isogenic RPE clones with stable mtDNA heteroplasmy from 3% to 99% for a Scal-induced mitochondrial deletion (mtDNA^ΔScal^) was chosen for study^29^. The reported heteroplasmy levels of these clones were reconfirmed by droplet digital PCR (Fig. 1a). The Scal-deleted region of mtDNA includes seven protein-encoding genes and three tRNAs (Extended Data Fig. 1a).

**Fig. 1:**
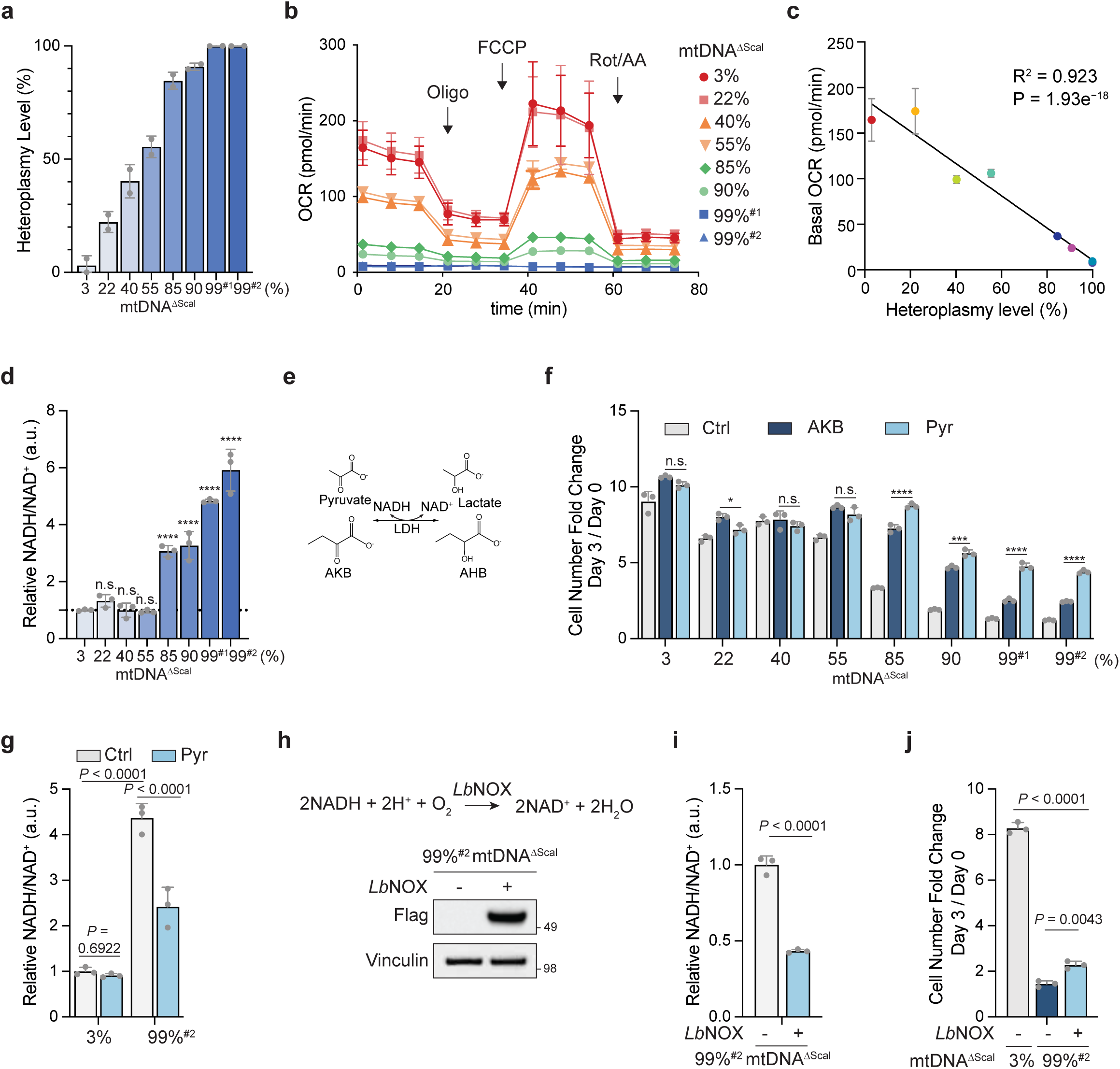
mtDNA heteroplasmic cells exhibit defective oxidative metabolism and impair proliferation. a, Droplet digital PCR (ddPCR) analysis of the levels of mtDNA^ΔScal^ heteroplasmy in RPE isolated single-cell clones. Bar plots are means ± s.d. from n = 2 replicates. b, Oxygen consumption rate (OCR) of indicated mtDNA heteroplasmy clones measured using Seahorse Bioanalyzer. Oligo, oligomycin; FCCP, carbonyl cyanide-p-trifluoromethoxyphenylhydrazone; Rot/AA, rotenone/antimycin A. Values represent mean ± s.d. from n = 4 replicates. c, Basal OCR (time point = 1 min) in (b) at indicated heteroplasmy levels. A simple linear regression between OCR and heteroplasmy levels is shown. For correlation analysis, the R^2^ and p-value are shown. d, Relative NADH/NAD^+^ ratio in the indicated mtDNA heteroplasmy clones cultured in uridine-supplemented DMEM without added pyruvate for 24 hours (h). The ratios were normalized to 3% mtDNA^ΔScal^ heteroplasmy. e, Schematic of the cytosolic LDH-catalyzed reduction of pyruvate and alpha-ketobutyrate (AKB) to lactate and alpha-hydroxybutyrate (AHB). f, Proliferation of indicated cells cultured with either 1 mM AKB or 1 mM pyruvate was measured by cell number fold change at day 3 relative to day 0. g, Relative NADH/NAD^+^ ratio in mtDNA heteroplasmy clones cultured in pyruvate-deficient DMEM with 1 mM pyruvate supplementation as indicated for 24 h. The ratios were normalized to 3% mtDNA^ΔScal^ heteroplasmy under control condition. h, Schematic of the reaction catalyzed by *Lb*NOX (top) and confirmation of *Lb*NOX expression in 99%^#2^ mtDNA^ΔScal^ cells by western blot (bottom). *Lb*NOX expression is detected using an anti-Flag tag antibody. Vinculin is used as a loading control. *Lb*NOX expression was induced by doxycycline (200 ng/mL) for 24 h. i, Relative NADH/NAD^+^ ratio in 99%^#2^ mtDNA^ΔScal^ cells with *Lb*NOX expression as indicated. The NADH/NAD^+^ ratios were normalized to conditions without doxycycline (200 ng/mL) induction. j, Proliferation of indicated cells with or without *Lb*NOX expression. Cell number fold change was measured at day 3 relative to day 0. 200 µM uridine was supplemented in all media unless otherwise indicated. Data are shown as mean ± s.d. from n = 3 independent replicates unless otherwise noted. Statistical significance was assessed using two-tailed t-tests (i), one-way ANOVA followed by Dunnett’s multiple comparisons test (d) or Tukey’s multiple comparisons test (f, j), two-way ANOVA (g),. ∗∗∗∗p < 0.0001; ∗∗∗p < 0.001; ∗∗p < 0.01; ∗p < 0.05; ns, nonsignificant.

The impact of mtDNA^ΔScal^ heteroplasmy on mitochondrial respiration was assessed using a Seahorse XFe96 Analyzer (Fig. 1b). Both basal and maximal oxygen consumption rates of the clones declined as the ratio of mtDNA^ΔScal^ to total mtDNA increased (Fig. 1c and Extended Data Fig. 1b), displaying a near-linear relationship (R^2^ = 0.923, *p* = 1.93e^-18^ for basal cellular oxygen consumption and R^2^ = 0.893, *p* = 3.07e^-16^ for maximal oxygen consumption). Previous reports have linked mtDNA heteroplasmy to impaired NADH oxidation due to defective electron transport chain (ETC) function. However, we found that the cellular NADH/NAD⁺ ratio remained unchanged in clones with ≤ 55% heteroplasmy. A significant increase in the cellular NADH/NAD^+^ ratio emerged only in the clones that had 85% or greater heteroplasmy (Fig. 1d). Levels of the electron transport chain components also remained stable in clones with 55% heteroplasmy or less. At higher levels of heteroplasmy, proteins in Complexes I to IV were reduced while the Complex V-associated protein ATP5A remained constant (Extended Data Fig. 1c). Cells with high heteroplasmy also exhibited reduced levels of the TCA metabolites citrate and malate and tracing of [U-^13^C] glucose revealed a decreased amount of glucose-derived pyruvate being oxidized in the TCA cycle, as measured by the (m+2) fraction of citrate and malate (Extended Data Fig. 1d-f).

The elevated NADH/NAD^+^ ratio observed in cells with 85% or greater heteroplasmy correlated with reduced proliferation despite the presence of 200 µM uridine in the medium (Fig. 1d). Previous studies have shown that exogenous electron acceptors such as pyruvate and alpha-ketobutyrate (AKB) can restore redox balance and support cell proliferation in respiration-compromised cells^15,26,27^ (Fig. 1e). In clones with ≤ 55% heteroplasmy, 1 mM pyruvate or 1 mM AKB had only modest effects on proliferation. In contrast, cells with 85% heteroplasmy more than doubled their growth and those with ≥ 90% heteroplasmy showed almost 3-fold increase in response to pyruvate. AKB supplementation also improved proliferation, but to a lesser extent (Fig. 1f). Consistent with its reported ability to buffer reductive stress, pyruvate decreased the NADH/NAD^+^ ratio in high-heteroplasmic cells to a significant extent (*p* < 0.0001) but not in cells with low heteroplasmy (*p* = 0.6922) (Fig. 1g).

To independently test whether NADH/NAD^+^ reduction is responsible for the restoration of proliferation, we introduced an inducible water-forming NADH oxidase from *Lactobacillus brevis* (*Lb*NOX) into cells with 99% heteroplasmy (Fig. 1h). Expression of *Lb*NOX reduced NADH/NAD^+^ ratio to a comparable extent as pyruvate (Fig. 1i). However, expression of *Lb*NOX only modestly increased proliferation (Fig. 1j).

### Amino acid supplementation enhances proliferation of high heteroplasmic cells

In addition to lowering the NADH/NAD^+^ ratio, treatment of high-heteroplasmic mtDNA^ΔScal^ cells with pyruvate also increased cellular citrate and malate levels (Extended Data Fig. 1g)^7^. Mitochondria can also contribute to non-essential amino acid biosynthesis and DMEM lacks five nonessential amino acids (NEAAs). Therefore, we also tested whether NEAAs supplementation could further enhance the proliferation of heteroplasmic cells in medium supplemented with 1 mM pyruvate and 200 µM uridine (Fig. 2 and Extended Data Fig. 2). NEAAs addition exhibited no effect on proliferation in clones with ≤ 85% heteroplasmy but significantly boosted proliferation in clones with 90% heteroplasmy, with even greater effects observed in 99% heteroplasmic cells (Fig. 2a). Similar results were observed when AKB was used in place of pyruvate (Fig. 2b). Uridine supplementation at 200 µM was present in this experiment as well and was utilized in the medium in all subsequent experiments unless otherwise noted. To rule out clonal variability, four additional isogenic clones with heteroplasmy levels of 88%, 92%^#1^, 92%^#2^, and 97% were cultured in either pyruvate- or AKB-containing medium. NEAAs supplementation significantly enhanced proliferation in each of these clones as well (Extended Data Fig. 2a).

**Fig. 2:**
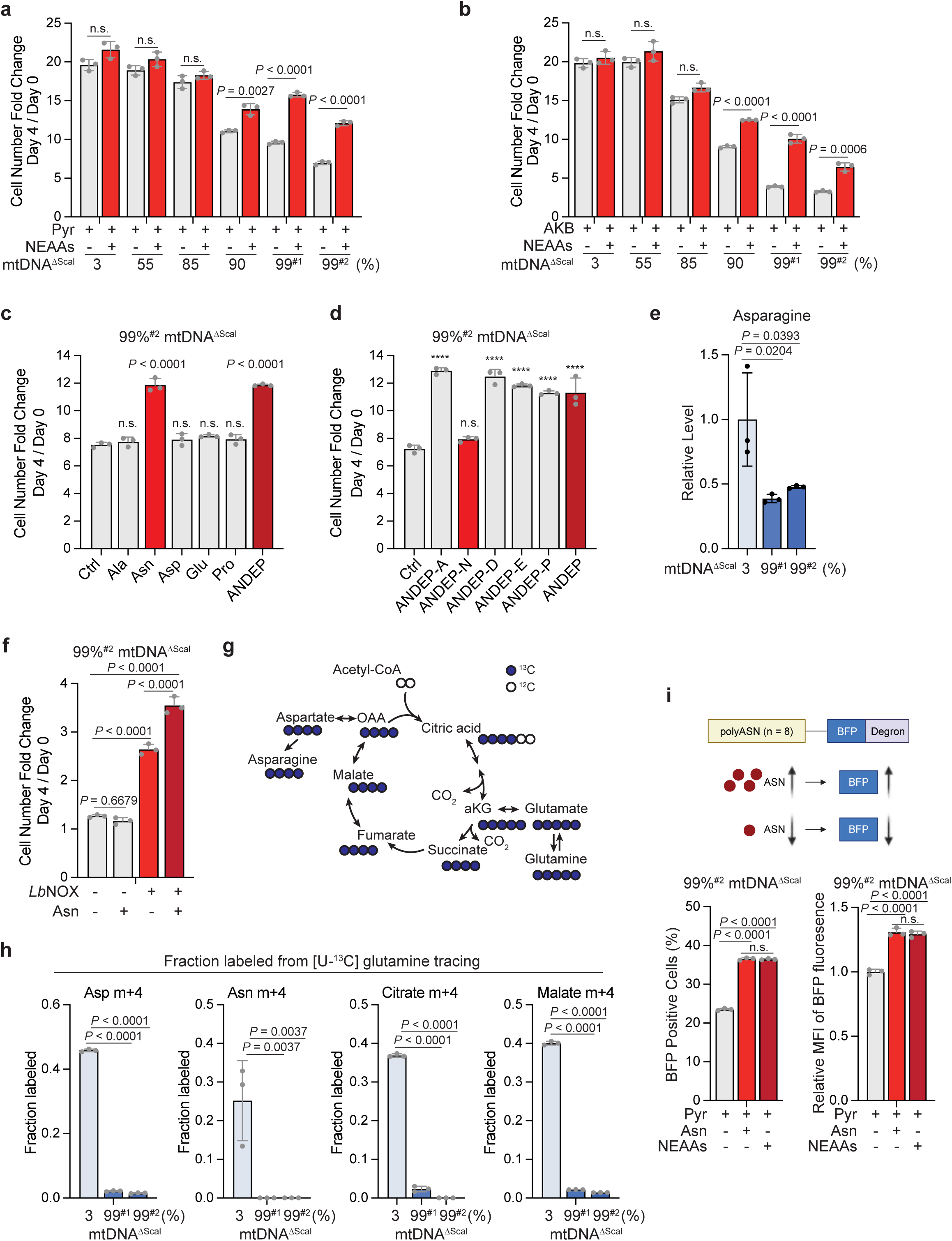
Asparagine supplementation enhances proliferation of high heteroplasmic cells. a,b, mtDNA heteroplasmic cells were cultured in DMEM supplemented with 1 mM pyruvate (a) or AKB (b), with 100 µM NEAAs as indicated. Pyr, pyruvate. Cell proliferation measured as cell number fold change (day 4/day 0). c, 99%^#2^ mtDNA^ΔScal^ cells were cultured in DMEM supplemented with 1 mM pyruvate, along with indicated non-essential amino acids. Ala, alanine; Asn, asparagine; Asp, aspartate; Glu, glutamate; Pro, proline. ANDEP, a mix of the individual amino acids. Each individual amino acid was supplemented at 100 µM. Cell proliferation measured as cell number fold change (day 4/day 0). d, Cells as in (c) were cultured in DMEM supplemented with 1 mM pyruvate along with addition of ANDEP lacking the indicated amino acids. A, alanine; N, asparagine; D, aspartate; E, glutamate; P, proline. ANDEP, a mix of the individual amino acids. e, Intracellular levels of asparagine in indicated mtDNA^ΔScal^ cells cultured in DMEM with 1 mM pyruvate for 8 h, measured by LC-MS. Relative metabolite levels were normalized to 3% mtDNA^ΔScal^ cells. f, Cell proliferation measured as cell number fold change (day 4/day 0) in 99%^#2^ mtDNA^ΔScal^ cells with *Lb*NOX expression and with 100 µM asparagine supplementation as indicated. Asn, asparagine. g, Schematic diagram of [U-^13^C] glutamine tracing. h, Fractional labeling of aspartate, asparagine, citrate, and malate pools by m+4 labeling in indicated mtDNA^ΔScal^ cells cultured in [U-^13^C] glutamine for 8 h, measured by LC-MS. 1 mM pyruvate was present in the medium. Asp, aspartate; Asn, asparagine. i, Schematic illustrating the design of the asparagine reporter (top). polyASN (n = 8), poly-asparagine tract expressing 8 asparagines; BFP, blue fluorescent protein. Percentage of BFP-positive cells (bottom left) and median fluorescence intensity (MFI, bottom right) was measured by FACS. 99%^#2^ mtDNA^ΔScal^ cells were transduced with an inducible polyASN-BFP construct in a retroviral vector. Cells were treated with DMEM containing 1 mM pyruvate with 100 µM asparagine or 100 µM NEAAs as indicated for 24 hr in the presence of doxycycline (200 ng/mL). Asn, asparagine. 200 µM uridine was supplemented in all media unless otherwise indicated. Data are shown as mean ± s.d. from n = 3 independent replicates. Statistical significance was assessed using two-tailed t-tests (a, b) or one-way ANOVA followed by Dunnett’s multiple comparisons test (c, d, e, h) or Tukey’s multiple comparisons test (f, i). ∗∗∗∗p < 0.0001; ns, nonsignificant.

To identify which of the five amino acids (alanine, asparagine, aspartate, glutamate, or proline), that are absent in DMEM but present in the amino acid supplement were responsible for the enhanced proliferation in high heteroplasmy cells, we added each amino acid individually to 99% heteroplasmic cells cultured in pyruvate- and uridine-containing medium. Only asparagine significantly enhanced proliferation, matching the effect of the full NEAAs supplement (Fig. 2c). Removal of asparagine from the NEAAs mix eliminated the ability of the NEAAs mix to promote proliferation (Fig. 2d). Metabolite profiling revealed that alanine, glutamate, and proline were elevated in high heteroplasmy cells. In contrast, both asparagine and aspartate were significantly reduced (Fig. 2e and Extended Data Fig. 2b). Although both aspartate and asparagine levels are significantly reduced in respiratory-compromised cells, only asparagine and not aspartate or the other three nonessential amino acids enhanced proliferation in 99% heteroplasmic cells cultured in pyruvate-supplemented medium (Fig. 2c). A similar result was observed, when pyruvate was replaced with AKB, asparagine enhanced proliferation but the other four amino acids did not (Extended Data Fig. 2c). To assess dose dependency, asparagine was supplemented across a range of concentrations (10-1000 µM). The ability of asparagine to enhance proliferation plateaued at asparagine levels ≥ 50 µM, a dose within the physiological range observed in human plasma^30^ (Extended Data Fig. 2d).

To independently validate the results, we tested 99% heteroplasmic cells with doxycycline-inducible *Lb*NOX expression. Asparagine supplementation alone did not significantly promote proliferation in the absence of doxycycline induction. However, when *Lb*NOX was induced, the high heteroplasmic cells resumed proliferative expansion and their expansion was significantly enhanced by the addition of 100 µM asparagine to the medium. (Fig. 2f and Extended Data Fig. 2e).

It has previously been reported that mammalian cancers depend on mitochondrial respiratory activity to maintain the production of aspartate required to engage in nucleotide production and cell proliferation^26,27^. However, we found that the addition of a physiologic level (100 µM) of aspartate to the medium failed to restore cellular proliferation. Furthermore, while we found cells with high heteroplasmy had significantly reduced levels of both aspartate and asparagine, levels of both amino acids remained measurable (Fig. 2e and Extended Data Fig. 2b). In these prior studies, cell proliferation of respiratory defective cancer cells could be rescued by supraphysiologic levels of aspartate or by overexpressing cellular aspartate transporter SLC1A3^31,32^. Whether nontransformed RPE cells, which have comparable levels of respiration compromise can be similarly rescued by such manipulations is uncertain. Therefore, we tested whether either of these manipulations could similarly rescue the proliferation of RPE cells with 99% heteroplasmy for mtDNA^ΔScal^ cultured in uridine supplemented medium without pyruvate. As previously published doses of aspartate in excess of 2 mM could restore substantial proliferation of respiratory-deficient RPE cells even in the absence of supplemental pyruvate (Extended Data Fig. 2f). Similarly, overexpression of SLC1A3 in RPE cells led to the ability of these cells but not control cells, to proliferate at physiologic levels of aspartate 100 µM. In neither the present studies (Fig.2) or prior studies, was aspartate found to enhance the proliferation of cells grown in medium supplemented with pyruvate, suggesting that supplemental aspartate at a physiological concentration had no further effect on protein and nucleotide biosynthesis in respiratory compromised cells when pyruvate is present at sufficient levels to reduce reductive stress. But our data suggested that at least RPE cells depended on exogenous asparagine for optimal cell growth even when pyruvate supplementation is provided.

In respiratory-competent cells, glutamine carbon supports the production of asparagine (Fig. 2g). Using isotope tracing with [U-^13^C] glutamine, we found that asparagine (m+4) produced from glutamine labeled 25% of the asparagine pool at 8 hours in cells with 3% heteroplasmy. In contrast, no asparagine synthesis from glutamine entering the oxidative TCA cycle was observed over the same time scale in cells with 99% mtDNA heteroplasmy. The contribution of glutamine to aspartate (m+4), citrate (m+4), and malate (m+4) was also compromised in high heteroplasmic clones confirming that the cells lacked significant TCA cycle activity (Fig. 2h). However, glutamine-dependent reductive carboxylation did contribute significantly more to the remaining pools of citrate, aspartate, asparagine in respiratory compromised cells with 99% mutant mtDNA heteroplasmy than it did to cells with 3% mutant mtDNA heteroplasmy consistent with prior studies^33^ (Extended Data Fig. 2g-h).

Unlike aspartate, which is essential for both de novo nucleotide synthesis and protein synthesis, asparagine is used exclusively for protein synthesis^34^. Therefore, if the reduced level of asparagine observed in cells with high mtDNA^ΔScal^ heteroplasmy is biologically significant, there should be a reduction in mRNA translation in the absence of asparagine supplementation. Consistent with this, we found that while pyruvate addition to uridine-supplemented DMEM increased the bulk translation rate of RPE cells with 99% mtDNA^ΔScal^ heteroplasmy, supplementation with 100 µM asparagine but not 100 µM aspartate led to an additional enhancement of bulk protein production across a broad size range when measured through incorporation of puromycin, a tyrosyl-tRNA mimetic, into total cellular proteins by Western blotting (Extended Data Fig. 2i). To more rigorously test that the increase in bulk protein expression is due to enhanced asparagine availability, we developed a translation-based asparagine reporter using an inducible polyASN protein reporter (PolyASN-BFP) comprising an 8-residue poly-asparagine tract fused via a flexible linker to a blue fluorescent protein open reading frame (BFP ORF). A degron from mouse ornithine decarboxylase was added to the BFP ORF to reduce the half-life of the BFP reporter^35,36^ (Fig. 2i). The reporter was transduced into 99% heteroplasmic cells. In 99% heteroplasmic cells, supplementation with asparagine or NEAAs increased both the proportion of BFP-positive cells and their mean fluorescence intensity (MFI) (Fig. 2i).

### Asparagine limitation activates the integrated stress response and inhibits protein synthesis

To examine how the proteome of cells with high heteroplasmic mtDNA^ΔScal^ levels is affected by exogenous asparagine, we performed untargeted proteomic profiling in cells with 99% heteroplasmy cultured in pyruvate- and uridine-supplemented medium in the presence or absence of exogenous asparagine. Comparison revealed 22 proteins significantly upregulated and 32 downregulated in asparagine-deficient conditions. Notably, among the upregulated proteins, seven were known ATF4 targets (Fig. 3a). Gene Set Enrichment Analysis (GSEA) demonstrated a significant enrichment of the Reactome ATF4 pathway gene set in the absence of exogenous asparagine (NES = 1.71, *p*-adj = 0.0489) (Fig. 3b). Conversely, the epithelial-to-mesenchymal-transition (EMT) pathway was the most downregulated pathway in asparagine-deficient medium (Extended Data Fig. 3a). One of the known ATF4 target genes that was upregulated is PCK2, which encodes the enzyme mitochondrial phosphoenolpyruvate carboxykinase (PEPCK-M) that catalyzes the conversion of oxaloacetate (OAA), the carbon backbone of both aspartate and asparagine, to phosphoenolpyruvate (PEP). To test whether PEPCK-M-dependent diversion of OAA into PEP limits aspartate and asparagine production, we generated PCK2-knockout cells using CRISPR-Cas9 in RPE cells with 99%^#2^ heteroplasmy mtDNA^ΔScal^. Loss of PCK2 partially reduced rather than promoted the proliferation of the high heteroplasmic cells when grown in pyruvate- and uridine-supplemented medium. Furthermore, the addition of asparagine still significantly enhanced the proliferation of the high heteroplasmic cells deficient in PCK2 (Extended Data Fig. 3b).

**Fig. 3:**
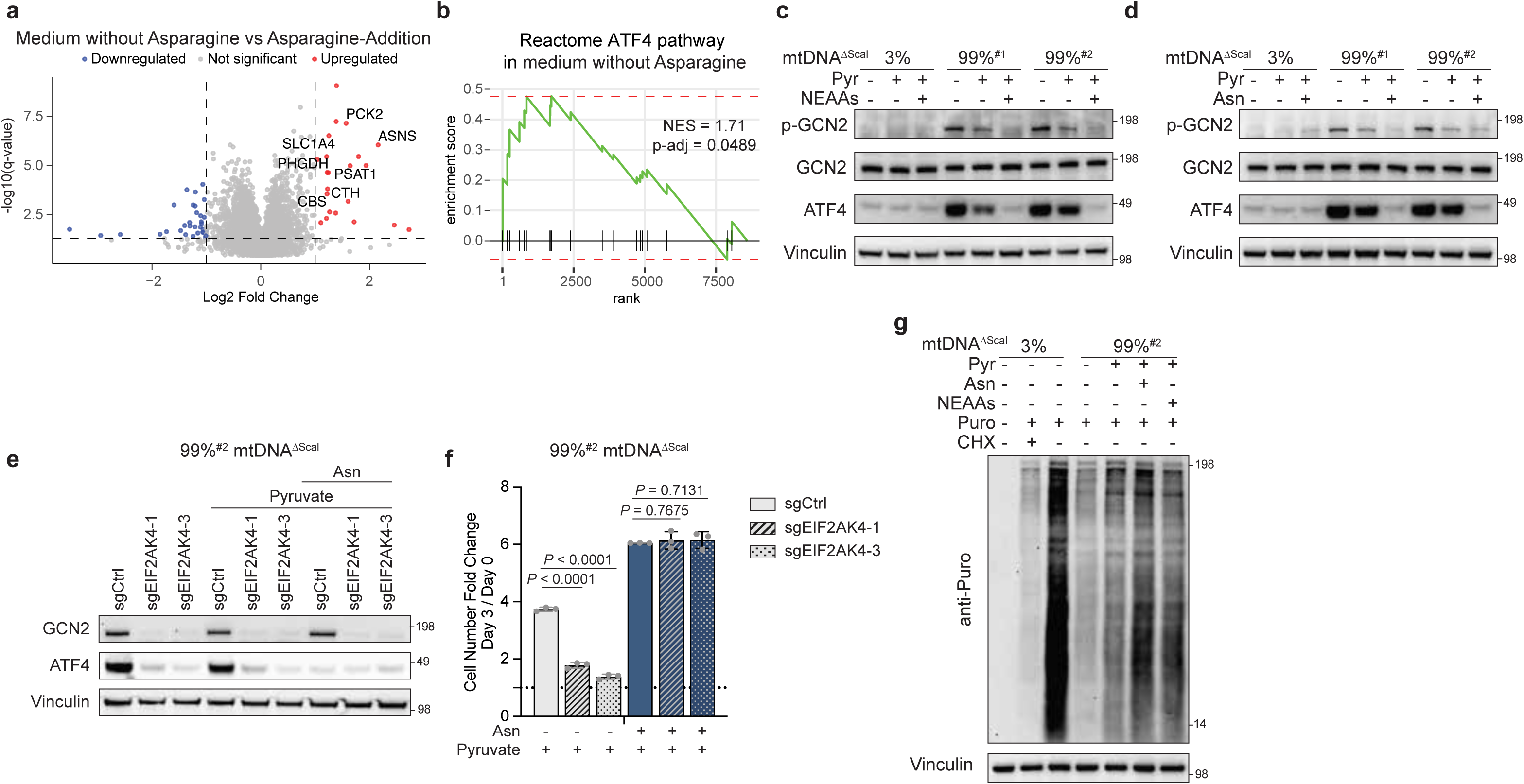
Asparagine limitation activates integrated stress response and inhibits protein translation. a, Volcano plots representing the proteome analysis of proteins in 99%^#2^ mtDNA^ΔScal^ cells cultured in DMEM containing 1 mM pyruvate with or without 100 µM asparagine as indicated. Highlighted points (red and blue) are statistically significant (q-value < 0.05) and fold change ≥ 2. Gene names are given to points that are ATF4 targets as per the Reactome ATF4 pathway. b, GSEA of the proteomics data showing the upregulation of the Reactome ATF4 pathway in 99%^#2^ mtDNA^ΔScal^ cells cultured in DMEM without asparagine supplementation. c,d, mtDNA heteroplasmic cells were treated with 1 mM pyruvate and 100 µM NEAAs (c) or 100 µM asparagine (d) as indicated for 24 h, followed by immunoblotting with the indicated antibodies. e,f, Western blot (e) and cell proliferation (f) of 99%^#2^ mtDNA^ΔScal^ cells with sgCtrl (sgPRM1), sgEIF2AK4 (GCN2)-1 or sgEIF2AK4 (GCN2)-3 mediated CRISPR-Cas9 genome editing cultured in the presence of 1 mM pyruvate and 100 µM asparagine supplementation as indicated. Cells were cultured in indicated medium for 24 h for (e). Data in (f) are shown as mean ± s.d. from n = 3 independent replicates. Statistical significance was assessed using two-way ANOVA followed by Dunnett’s multiple comparisons test. g, 3% and 99%^#2^ mtDNA^ΔScal^ cells were cultured in the indicated medium for 24 h. Puromycin (90 µM) was added 10 min before sample collection, followed by immunoblotting with puromycin antibody. CHX, cycloheximide, a general translation inhibitor, was used at 10 µg/mL 30 min prior to puromycin treatment as control. 200 µM uridine was supplemented in all media unless otherwise indicated.

To test for ATF4 induction in the high heteroplasmic clones, we examined ATF4 protein levels in two isogenic 99% mtDNA heteroplasmy clones cultured in DMEM supplemented with either pyruvate alone or pyruvate plus NEAAs addition. Uridine was included in all conditions unless otherwise indicated. Both 99%^#1^ and 99%^#2^ clones had elevated ATF4 expression in the absence of pyruvate and NEAAs. Pyruvate supplementation modestly downregulated ATF4 levels. Further addition of NEAAs or asparagine suppressed ATF4 to near baseline levels (Fig. 3c and 3d). Either asparagine or NEAAs supplementation also decreased ATF4 protein levels when high heteroplasmic cells were cultured in the presence of AKB (Extended Data Fig. 3c). In contrast, the isogenic clone with 3% heteroplasmy expressed significantly less ATF4 and the ATF4 level was unchanged by supplementation with pyruvate or pyruvate plus asparagine (Fig. 3c and 3d). Consistent with the role of ATF4 as a transcription factor, mRNA levels of ATF4 target genes were reduced upon asparagine supplementation in 99%^#2^ clones (Extended Data Fig. 3d).

ATF4 is a central effector of the integrated stress response (ISR), often activated through phosphorylation of eIF2α by stress-responsive kinases^37^. Since asparagine is exclusively used for tRNA charging and translation^34^, we hypothesized that uncharged tRNA^Asn^ might lead to autophosphorylation of general control non-derepressible 2 (GCN2) kinase. GCN2 was expressed at comparable levels in both the low and high heteroplasmy clones (Fig. 3c and 3d). However, GCN2 phosphorylation was only observed in the high heteroplasmy clones. Pyruvate supplementation partially reduced GCN2 phosphorylation. When both pyruvate and supplemental NEAAs mixtures were added, GCN2 phosphorylation was reduced to the levels observed in low-heteroplasmy cells (Fig. 3c). A comparable effect on GCN2 phosphorylation was observed when only asparagine was added along with pyruvate (Fig. 3d).

To confirm that GCN2 is required to activate ATF4 in response to asparagine deficiency, we used CRISPR-Cas9 to knock out the GCN2-encoding gene *EIF2AK4* in cells with 99% heteroplasmy for mtDNA^ΔScal^. *EIF2AK4* knockout was sufficient to abolish ATF4 activation irrespective of pyruvate or asparagine supplementation (Fig. 3e). High heteroplasmic cells with *EIF2AK4* deleted failed to upregulate ATF4 and exhibited a more pronounced growth defect in pyruvate- and uridine-containing medium (Fig. 3f). Asparagine supplementation was sufficient to rescue cell proliferation in both the control and GCN2-deleted cells (Fig. 3f). In contrast, knockout of heme-regulated eIF2α kinase (HRI)-encoding gene *EIF2AK1*, the other kinase implicated in ATF4 induction through mitochondrial dysfunction, exhibited no influence on ATF4 activation or cellular proliferation in 99% heteroplasmic cells (Extended Data Fig. 3e and 3f).

GCN2 phosphorylation is associated not only with the increased levels of proteins involved in the ISR but also with generalized repression of global translation. To test if exogenous asparagine addition by suppressing GCN2 phosphorylation led to an increase in total protein translation, we measured the effect of either exogenous asparagine or NEAAs on incorporation of puromycin (Fig. 3g and Extended Data Fig. 3g). Both conditions led to a reproducible increase in incorporated puromycin into cellular proteins.

### High heteroplasmy cells can utilize pyruvate carboxylase to maintain asparagine synthesis

In respiration competent cells, glucose and glutamine are both potential contributors to the production of oxaloacetate, the carbon backbone of both aspartate and asparagine. With the absence of TCA cycle-driven respiration, glutamine can still be used to produce oxaloacetate through reductive carboxylation. The oxaloacetate produced occurs in the cytoplasm where ATP-citrate lyase (ACLY) converts citrate to oxaloacetate and acetyl-CoA. In contrast, pyruvate carboxylase (PC) is a mitochondrial protein that carboxylates glucose-derived pyruvate into oxaloacetate. Therefore, we tested whether PC could potentially contribute to the generation of asparagine in respiratory-compromised cells^38–41^. Using isotope labeled tracing, we assessed PC activity by measuring the generation of asparagine (m+3) from [U-^13^C] glucose-derived pyruvate (m+3), as diagrammed in Fig. 4a. As a positive control we also analyzed 143B *cytochrome b* cybrid (143B cytB) cells, whose mitochondria have a hypomorphic mutation in the mtDNA encoded *cytochrome b* (*MT-CYB*) gene that reduced respiratory capacity^28^. 143B cytB cells are not reported to be asparagine-deficient^31^ and we found that the 143B cytB cells had an almost 6-fold higher levels of asparagine than RPE cells with 99% heteroplasmy for mtDNA^ΔScal^ (Extended Data Fig. 4a). We found that the amount of labeling asparagine (m+3) generated from [U-^13^C] glucose through PC at 6 hours was more than 10-fold higher in 143B cytB cells than in RPE cells with 99% heteroplasmy for mtDNA^ΔScal^ (Fig. 4b). Western blot analysis of PC expression revealed that the difference in asparagine labeling correlated with a substantial difference in PC protein levels between the two cell lines (Fig. 4c). PC-dependent flux of [U-^13^C] glucose to citrate (m+5), malate (m+3) and aspartate (m+3) were also greater in 143B cytB cells in comparison to RPE 99% mtDNA^ΔScal^ cells (Extended Data Fig. 4b).

**Fig. 4:**
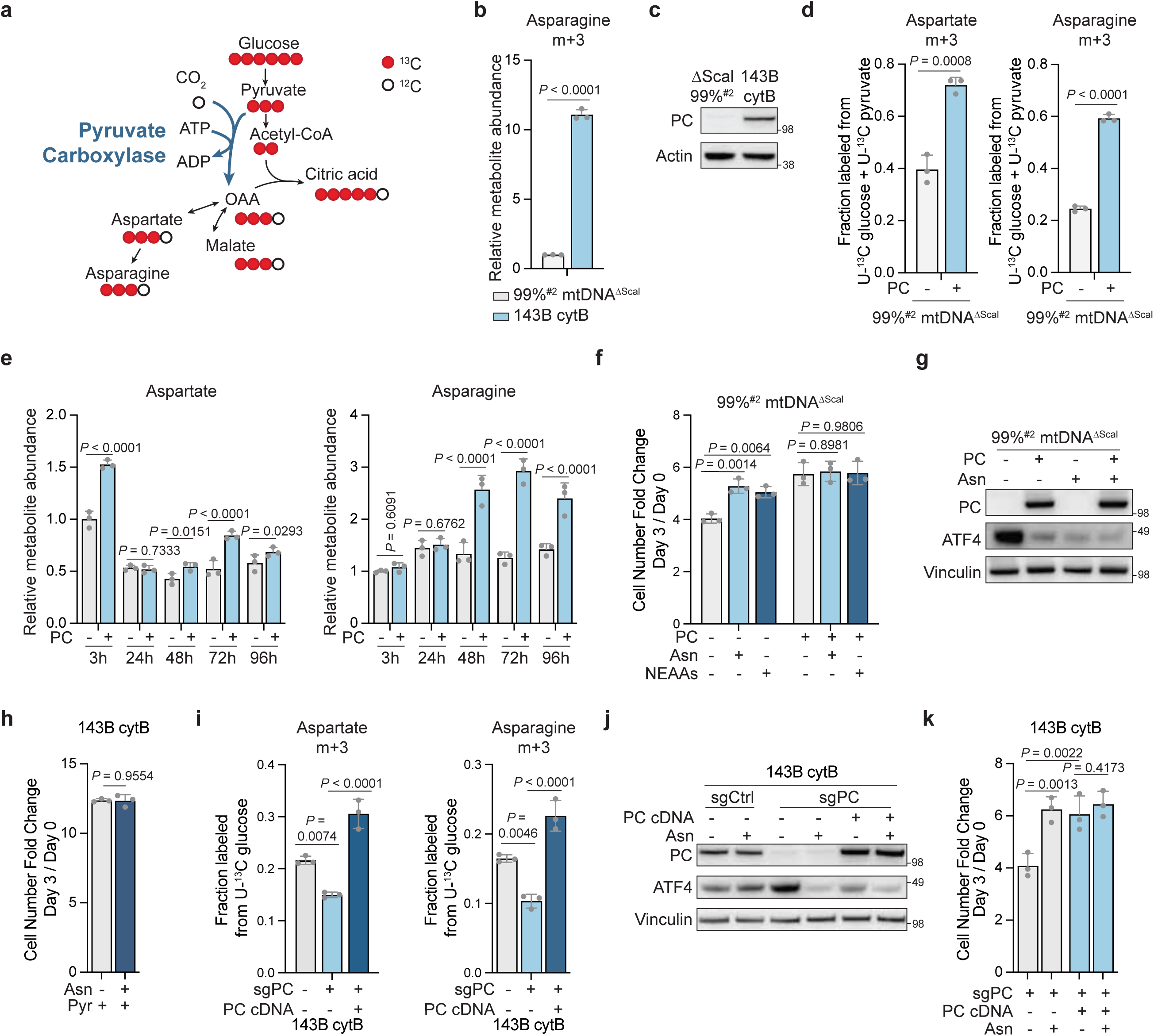
High heteroplasmy cells are dependent on pyruvate carboxylase for asparagine synthesis. a, Schematic diagram of the [U-^13^C] glucose tracing experiment. b, Relative metabolite abundance of isotope-labeled asparagine (m+3) measured by LC-MS in the indicated cells. Cells were cultured in glucose-deficient DMEM containing 10 mM [U-^13^C] glucose and 1 mM AKB for 6 h. c, Immunoblot of PC expression levels in 99%^#2^ mtDNA^ΔScal^ (RPE) and 143B cytB cells. d, Fractional labeling of aspartate (m+3) and asparagine (m+3) in 99%^#2^ mtDNA^ΔScal^ cells expressing empty vector control or PC cultured in glucose-deficient DMEM containing 10 mM [U-^13^C] glucose and 1 mM [U-^13^C] pyruvate for 24 h, measured by LC-MS. e, Relative metabolite abundance of aspartate and asparagine in 99%^#2^ mtDNA^ΔScal^ cells expressing empty vector control or PC cultured in DMEM with pyruvate supplementation for indicated time. f, Cell proliferation measured as cell number fold change (day 3/day 0) in 99%^#2^ mtDNA^ΔScal^ cells expressing empty vector control or PC cDNA cultured in medium with 100 µM asparagine or NEAAs as indicated and with 1 mM pyruvate present across all conditions. g, Western blot of 99%^#2^ mtDNA^ΔScal^ cells expressing empty vector control or PC cultured in 1 mM pyruvate with 100 µM asparagine supplementation as indicated. h, Cell proliferation measured as cell number fold change (day 3/day 0) in 143B cytB cells with or without 100 µM asparagine supplementation as indicated. 1 mM pyruvate was supplemented in both conditions. i, Fractional labeling of aspartate (m+3) and asparagine (m+3) in 143B cytB cells expressing sgCtrl or sgPC and with PC cDNA as indicated. sgPC (-) indicates the expression of sgPRM1 as a control. Cells were cultured in glucose-deficient DMEM containing 10 mM [U-^13^C] glucose and 1 mM AKB for 6 h, measured by LC-MS. j, Western blot of 143B cytB cells expressing sgCtrl or sgPC and with PC cDNA as indicated, cultured with 100 µM asparagine supplementation as indicated. 1 mM pyruvate was supplemented across all conditions. k, Cell proliferation measured as cell number fold change (day 3/day 0) in 143B cytB cells expressing sgPC with PC cDNA and cultured in medium supplemented with 100 µM asparagine as indicated. 1 mM pyruvate was supplemented across all conditions. 200 µM uridine was supplemented in all the media unless otherwise indicated. Data are shown as mean ± s.d. from n = 3 independent replicates. Statistical significance was assessed using two-tailed t-tests (b, d, h), one-way ANOVA followed by Tukey’s multiple comparisons test (i), two-way ANOVA (e, f, k) followed by Tukey’s multiple comparisons test (e) or Dunnett’s multiple comparisons test (f). Asn, asparagine. Pyr, pyruvate.

To explore the possibility that the relative PC deficiency enhanced RPE cells dependency on exogenous asparagine, an ectopic PC construct was stably introduced into clones with 99% mtDNA^ΔScal^ heteroplasmy. Since respiratory-deficient cells are normally cultured in pyruvate-supplemented medium, which can dilute the entry of [U-^13^C] glucose-derived pyruvate into the mitochondrial matrix, we used a combination of 10 mM [U-^13^C] glucose and 1 mM [U-^13^C] pyruvate tracing to assess the full extent to which PC-derived oxaloacetate contributed to aspartate and asparagine in respiratory-compromised cells. Isotope labeled [U-^13^C] glucose and [U-^13^C] pyruvate tracing showed that PC-derived aspartate (m+3) increased from 40% to 72%, and asparagine (m+3) increased from 25% to 59% in the 99% mtDNA^ΔScal^ RPE cells with PC introduction (Fig. 4d). Increased percentage labeling of PC-derived TCA metabolites citrate (m+5), malate (m+3), and fumarate (m+3) were also observed in the PC-expressing cells (Extended Data Fig. 4c).

To determine whether increased flux through PC elevated aspartate and asparagine levels, metabolite abundance was compared between PC-expressing and control 99% mtDNA^ΔScal^ cells following replacement of the medium in which they were passaged to DMEM supplemented with pyruvate and uridine but without additional NEAAs. The most significant difference observed in the levels of aspartate was that PC-expressing cells had higher levels at 3 hours after switching the medium; however, aspartate levels declined in both PC-expressing and control cells within the first 24 hours and there was only a modest elevation of aspartate levels in PC expressing cells at later time point. In contrast, total asparagine levels were similar in PC-expressing and control cells at 3 hours. However, total levels of asparagine increased in PC-expressing cells at subsequent time points, while there was little change in control cells over time. By 48-96 hours, asparagine levels were significantly higher in PC-expressing cells than in control cells, and the PC-expressing cells were proliferating more (Fig. 4e and 4f).

Ectopic PC expression enhanced proliferation to a greater extent than either the addition of NEAAs or asparagine and rendered the proliferation of high heteroplasmic cells independent of exogenous asparagine (Fig. 4f). When cells were cultured in uridine- and pyruvate-supplemented DMEM without NEAAs supplementation, PC expression was associated with reduced ATF4 activation in comparison to control cells. The degree of this suppression was comparable to direct asparagine supplementation (Fig. 4g). Similar restoration of proliferation and suppression of ATF4 was obtained in AKB- and uridine-containing medium (Extended Data Fig. 4d and 4e). PC expression also enhanced bulk cellular protein production to a level comparable to that achieved with asparagine supplementation (Extended Data Fig. 4f).

### Endogenous pyruvate carboxylase is required for 143B cytB cell proliferation in the absence of exogenous asparagine

143B cytB cells have been widely used as a model of mammalian cells with hypomorphic mtDNA^26,27,31^. Unlike RPE cells, these osteosarcoma cybrids express high PC protein levels and efficiently synthesize asparagine (m+3) from glucose (Fig. 4b, 4c, and Extended Data Fig. 4a, 4b). Consistently, exogenous asparagine did not significantly enhance proliferation of 143B cytB cells cultured in pyruvate- and uridine-containing medium (Fig. 4h), suggesting that endogenous PC activity is sufficient to support asparagine biosynthesis. To directly confirm this, we used CRISPR-Cas9 to delete PC in 143B cytB cells and performed [U-^13^C] glucose tracing. The aspartate (m+3) and asparagine (m+3) fraction generated from glucose was significantly decreased in 143B cytB cells when PC was deleted and was rescued to a level greater than wild-type (WT) when the PC-knockout cells were transduced with a CRISPR-Cas9-resistant PC cDNA (Fig. 4i). Malate (m+3), fumarate (m+3), and citrate (m+5) responded similarly to aspartate and asparagine following PC deletion and restoration (Extended Data Fig. 5a). PC deletion also activated ATF4 in 143B cytB cells that were maintained in pyruvate- and uridine-supplemented DMEM, and ATF4 expression was suppressed by either extracellular asparagine supplementation or by the reintroduction of PC (Fig. 4j). The PC-deleted 143B cytB cells exhibited reduced proliferation in DMEM supplemented with pyruvate and uridine, which could be enhanced by asparagine. Reintroduction of a PC cDNA into the PC-knockout 143B cytB cells enhanced proliferation and abolished the asparagine dependency (Fig. 4k). AKB supplementation in place of pyruvate exhibited comparable results in terms of both ATF4 activation and proliferation (Extended Data Fig. 5b and 5c). Thus, the ability of PC to promote the cell growth of respiratory-compromised cells in the absence of exogenous asparagine does not depend on the presence of supraphysiologic levels of pyruvate in the medium (Extended Data Fig. 4d for RPE cells with high heteroplasmic mtDNA mutations and Extended Data Fig. 5c for 143B cytB cells).

**Fig. 5:**
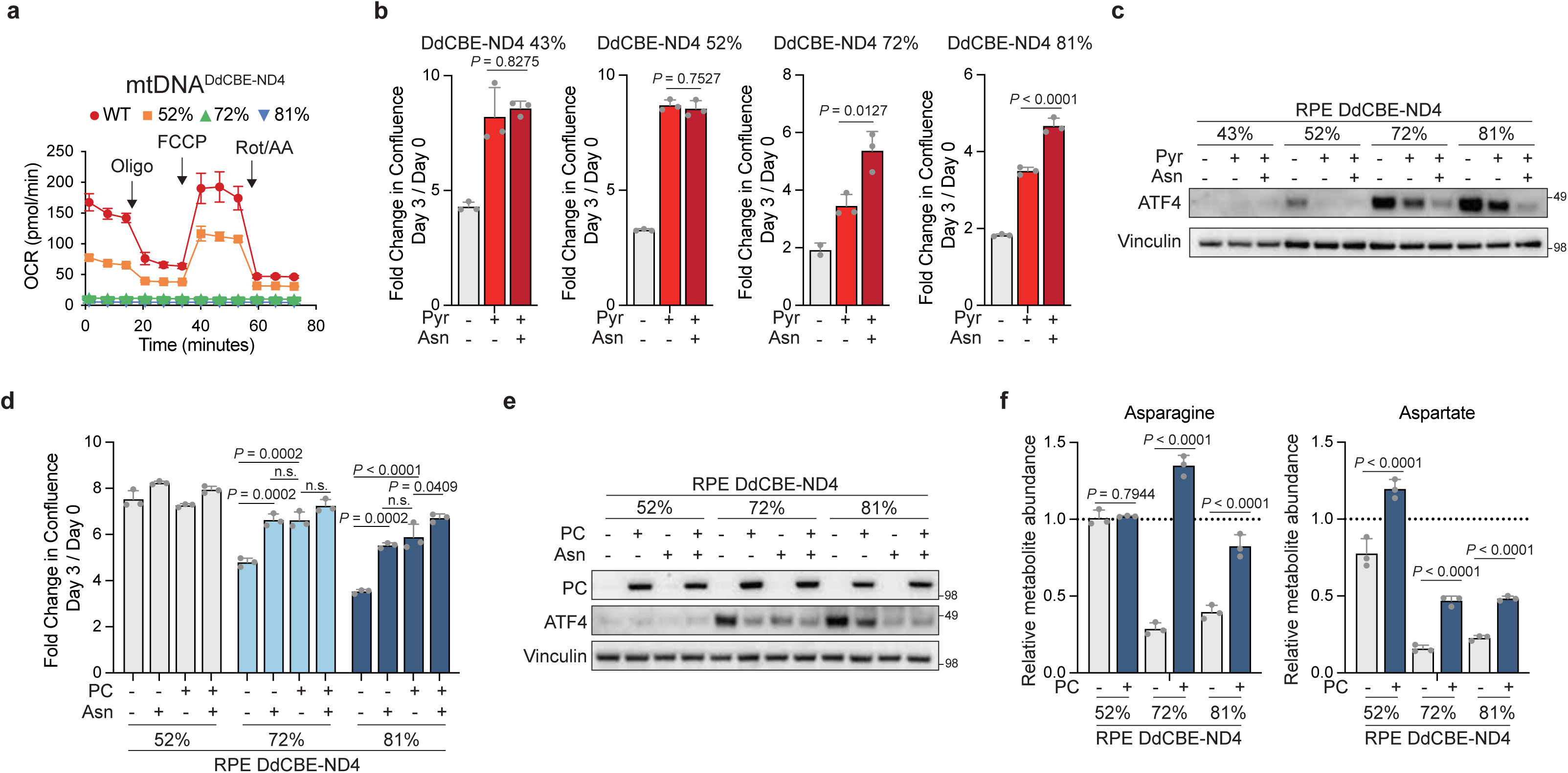
mtDNA point mutation cells exhibit heteroplasmy-dependent asparagine auxotrophy that is rescued by PC expression. a, Oxygen consumption rate (OCR) of RPE WT and indicated mtDNA^DdCBE-ND4^ heteroplasmy clones measured using Seahorse Bioanalyzer. Oligo, oligomycin; FCCP, carbonyl cyanide-p-trifluoromethoxyphenylhydrazone; Rot/AA, rotenone/antimycin A. Values represent mean ± s.d. from n = 6 replicates. b, Cell proliferation measured as fold change in confluence (day 3/day 0) of the indicated RPE mtDNA^DdCBE-ND4^ heteroplasmy clones, cultured in DMEM supplemented with 1 mM pyruvate or 100 µM asparagine as indicated. c, Western blot of the indicated RPE mtDNA^DdCBE-ND4^ heteroplasmy clones cultured in DMEM with 1 mM pyruvate or 100 µM asparagine supplementation as indicated. d, Cell proliferation measured as fold change in confluence (day 3/day 0) of the indicated RPE mtDNA^DdCBE-ND4^ heteroplasmy clones expressing empty vector control or PC cDNA, cultured in DMEM supplemented with 100 µM asparagine as indicated. 1 mM pyruvate was present across all conditions. e, Western blot of the indicated RPE mtDNA^DdCBE-ND4^ heteroplasmy clones expressing empty vector control or PC cDNA, cultured in DMEM with 100 µM asparagine supplementation as indicated. 1 mM pyruvate was present across all conditions. f, Relative metabolite abundance of asparagine and aspartate in the indicated RPE mtDNA^DdCBE-ND4^ heteroplasmy clones expressing empty vector control or PC cDNA, cultured in DMEM with 1 mM pyruvate supplementation for 24 h. Metabolite levels were normalized to those in RPE WT cells cultured under the same conditions, indicated by the dotted line. 200 µM uridine was supplemented across all conditions (b-f). Data are shown as mean ± s.d. from n = 3 independent replicates unless otherwise indicated. Statistical significance was assessed using one-way ANOVA followed by Tukey’s multiple comparisons test (b, d) or two-way ANOVA followed by Tukey’s multiple comparisons test (f).

### Cells with mtDNA point mutations exhibit heteroplasmy-dependent asparagine auxotrophy that is rescued by PC expression

To determine whether asparagine dependency in mtDNA high-heteroplasmy cells extends beyond the mtDNA deletion model, previously published DddA-derived cytosine base editors (DdCBEs) were used in RPE cells to catalyze a C•G-to-T•A conversion in mtDNA ND4, a component of Complex I^42–44^. A series of isogenic clones with heteroplasmy levels up to 81% (mtDNA^DdCBE-ND4^) was generated.

Cellular respiration was assessed by Seahorse Analyzer. The 52% mtDNA^DdCBE-ND4^ clone had reduced basal and maximal oxygen consumption rates compared with wild-type RPE cells, and the 72% and 81% mtDNA^DdCBE-ND4^ clones exhibited respiratory deficiency (Fig. 5a). Consistent with the mtDNA truncation model mtDNA^ΔScal^, mtDNA^DdCBE-ND4^ clones also showed decreased levels of TCA cycle metabolites citrate and malate, as well as aspartate and asparagine (Extended Data Fig. 6a). mtDNA^DdCBE-ND4^ clones with ≤ 52% heteroplasmy proliferated independently of exogenous asparagine in the presence of pyruvate and uridine. However, at heteroplasmy levels ≥ 72%, asparagine supplementation enhanced proliferation (Fig. 5b). In the presence of uridine, the ISR indicator ATF4 was upregulated in the 52% mtDNA^DdCBE-ND4^ clone but was suppressed by pyruvate supplementation. No ATF4 was induced in the 43% mtDNA^DdCBE-ND4^ clone grown in uridine- or uridine- and pyruvate-containing medium. In contrast, the 72% and 81% mtDNA^DdCBE-ND4^ clones grown in uridine-containing medium expressed high levels of ATF4 and pyruvate supplementation only modestly reduced ATF4 levels, whereas asparagine addition decreased ATF4 to baseline levels (Fig. 5c).

To test whether limited PC activity constrains asparagine synthesis in high-heteroplasmy clones, PC cDNA was introduced into the 52%, 72%, and 81% mtDNA^DdCBE-ND4^ clones. Ectopic PC expression did not significantly affect proliferation in the 52% mtDNA^DdCBE-ND4^ clone, but enhanced proliferation in the 72% and 81% mtDNA^DdCBE-ND4^ clones to levels of exogenous asparagine addition (Fig. 5d). PC expression also suppressed ATF4 levels without extracellular asparagine supplementation in the 72% and 81% mtDNA^DdCBE-ND4^ clones (Fig. 5e). PC introduction did not alter asparagine levels in 52% mtDNA^DdCBE-ND4^ clones but increased asparagine levels in the 72% and 81% mtDNA^DdCBE-ND4^ clones to levels comparable to wild-type RPE cells. Aspartate levels were also increased across all mtDNA^DdCBE-ND4^ clones but at a less extent than for asparagine (Fig. 5f). TCA cycle metabolites such as malate were also increased with PC expression (Extended Data Fig. 6b).

### Asparagine auxotrophy in respiration-deficient cells beyond mtDNA defects

To determine whether asparagine auxotrophy in respiratory-deficient cells is restricted to mtDNA heteroplasmy, we used CRISPR-Cas9 to knock out the nuclear-encoded ETC components *UQCRC2* (Complex III) and *COX4I1* (Complex IV) in RPE WT cells, the parental cell line for mtDNA^ΔScal^ and mtDNA^DdCBE-ND4^ clones (Fig. 6a). Cellular respiration was substantially reduced in *UQCRC2* or *COX4I1* knockout cells (Fig. 6b). Asparagine promoted proliferation in the presence of pyruvate and uridine to a similar extent as that observed in mtDNA^ΔScal^ and mtDNA^DdCBE-ND4^ high heteroplasmy cells (Fig. 6c). PC introduction enhanced proliferation independent of asparagine supplementation (Fig. 6d).

**Fig. 6:**
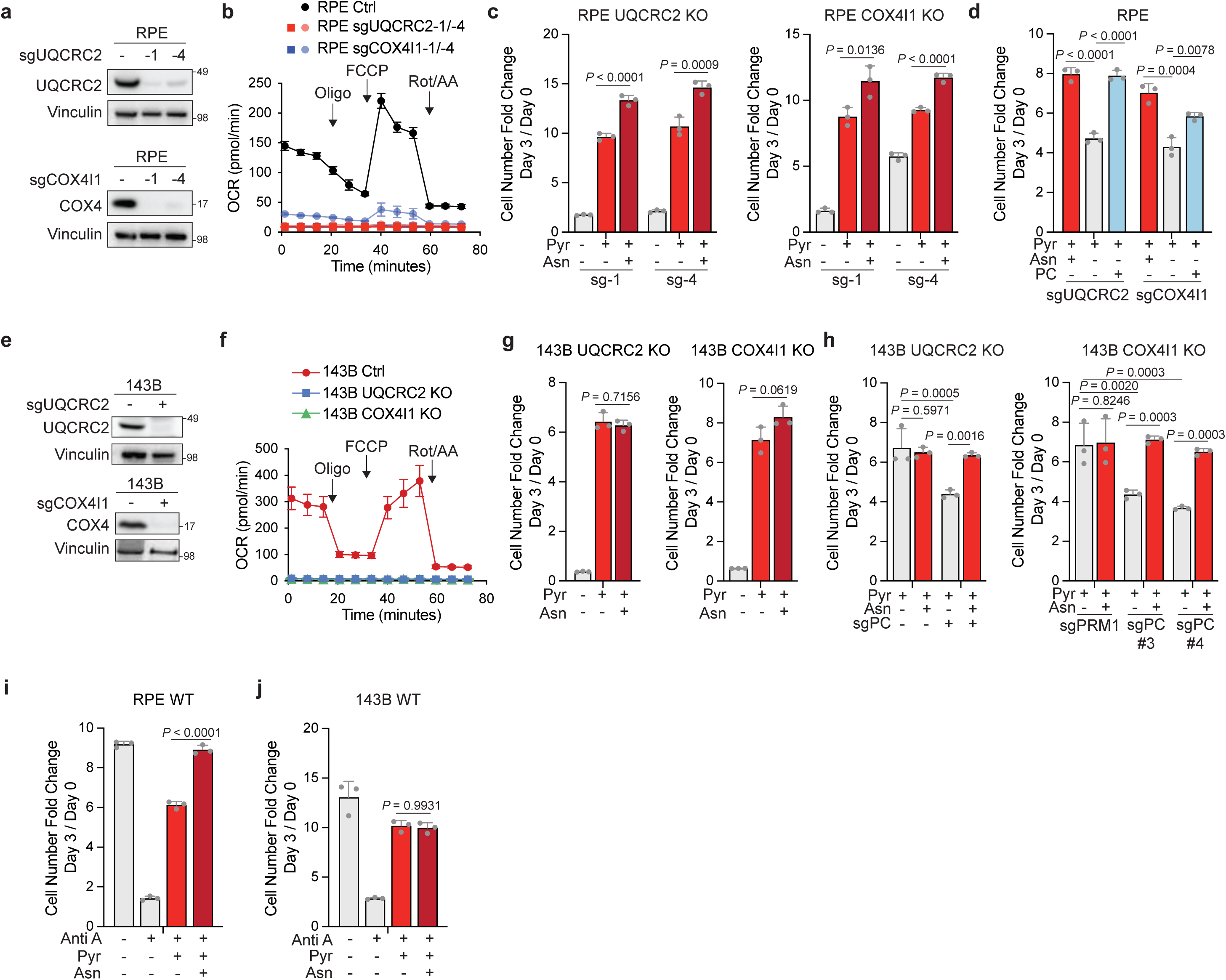
Asparagine auxotrophy in respiration-deficient cells beyond mtDNA defects. a-c, Western blot (a), oxygen consumption rate (b), and cell proliferation measured as cell number fold change (day 3/day 0) (c) of RPE WT cells with sgUQCRC2-1, sgUQCRC2-4, sgCOX4I1-1, or sgCOX4I1-4 mediated CRISPR-Cas9 genome editing, cultured in the presence of 1 mM pyruvate and 100 µM asparagine supplementation as indicated. sgUQCRC2 (-) or sgCOX4I1 (-) indicates the expression of sgPRM1 as a control. d, Cell proliferation measured as cell number fold change (day 3/day 0) in RPE UQCRC2 KO cells and COX4I1 KO cells expressing empty vector control or PC cDNA cultured in medium with 100 µM asparagine as indicated. 1 mM pyruvate present across all conditions. e-g, Western blot (e), oxygen consumption rate (f), and cell proliferation measured as cell number fold change (day 3/day 0) (g) of 143B WT cells with sgUQCRC2, or sgCOX4I1 mediated CRISPR-Cas9 genome editing, cultured in the presence of 1 mM pyruvate and 100 µM asparagine supplementation as indicated. sgUQCRC2 (-) or sgCOX4I1 (-) indicates the expression of sgPRM1 as a control. h, Cell proliferation measured as cell number fold change (day 3/day 0) in 143B UQCRC2 KO cells and COX4I1 KO cells expressing sgPC for CRISPR-Cas9 mediated PC KO, cultured in medium with 100 µM asparagine as indicated. 1 mM pyruvate present across all conditions. i-j, Cell proliferation measured as cell number fold change in RPE WT cells (i) and 143B WT cells (j) cultured in DMEM with 2 µM (i) or 100 nM (j) Antimycin A, 1 mM pyruvate, or 100 µM asparagine as indicated. 200 µM uridine was supplemented across all conditions unless otherwise indicated (a, c-e, g-j). Data are shown as mean ± s.d. from n = 3 independent replicates across all proliferation assays (c, d, g-j), n = 6 for Seahorse analysis in RPE cells (b), n = 5 in 143B cells (f). Statistical significance was assessed using one-way ANOVA followed by Tukey’s multiple comparisons test (c, d, g, i-j) and two-way ANOVA followed by Tukey’s multiple comparisons test (h).

In wildtype 143B cells, the parental line of 143B cytB cells, CRISPR-Cas9 deletion of *UQCRC2* or *COX4I1* also inhibited cellular respiration (Fig. 6e and 6f). Unlike RPE cells, asparagine did not affect the proliferation of 143B cells deleted of either *UQCRC2* or *COX4I1* when cultured in pyruvate- and uridine-supplemented medium (Fig. 6g). However, deletion of *PC* in either *UQCRC2* or *COX4I1* deficient 143B cells impaired proliferation, which could be significantly restored by addition of exogenous asparagine (Fig. 6h).

As an alternative way to inhibit cellular respiration, we also used the Complex III inhibitor Antimycin A to inhibit ETC activity. Cells treated with Antimycin A exhibited decreased cell proliferation in uridine-supplemented DMEM. This reduced proliferation could be partially rescued by pyruvate. Asparagine further promoted proliferation in RPE WT cells but had no effect on 143B WT cells (Fig. 6i and 6j).

### Tissue-specific PC levels regulate asparagine dependency under respiration inhibition

Tissue levels of PC vary widely, with high expression in liver and adipose tissue, whereas epithelial tissues display relatively low expression^45–48^. Despite this, several epithelial derived human cancers display a significant fraction of tumors with high heteroplasmic mtDNA mutations^49^. The most prevalent of these are thyroid, colorectal, and renal tumors^23–25^. To evaluate PC expression levels across cell lines, we compared three different thyroid cell lines and one colorectal cancer cell line with RPE and 143B cells. The thyroid cell lines include Nthy-ori 3-1, an immortalized normal thyroid cell line; NCI-237, a thyroid Hürthle cancer cell line, harboring a > 90% heteroplasmy for an mtDNA mutation in Complex I^50,51^; and TPC1, a papillary thyroid carcinoma cell line without known mtDNA mutation. All three thyroid lines exhibited lower PC expression than the 143B osteosarcoma line (Extended Data Fig. 7a).

Genetic knockout of ETC Complex III or Complex IV components in the normal thyroid cell line Nthy-ori 3-1 impaired cells proliferation even in the presence of pyruvate and uridine. Asparagine supplementation restored proliferation to the level comparable with WT cells (Extended Data Fig. 7b and 7c). Thus, nontransformed thyroid epithelial cells depend on ETC activity to suppress dependency on exogenous asparagine. To test whether cancer cell lines with low level PC expression also display dependency on exogenous asparagine, we tested TPC1 (thyroid cancer cells) and DLD1 (colorectal cancer cells) for the ability to proliferate in uridine-supplemented medium when treated with a respiratory inhibitor, either Complex I (Rotenone) or Complex III (Antimycin A). Rotenone or Antimycin A treatment reduced both TPC1 and DLD1 cell proliferation, which was partially rescued by pyruvate, and recovered to levels similar to untreated cells with both pyruvate and asparagine provided exogenously. (Extended Data Fig. 7d and 7e).

**Fig. 7:**
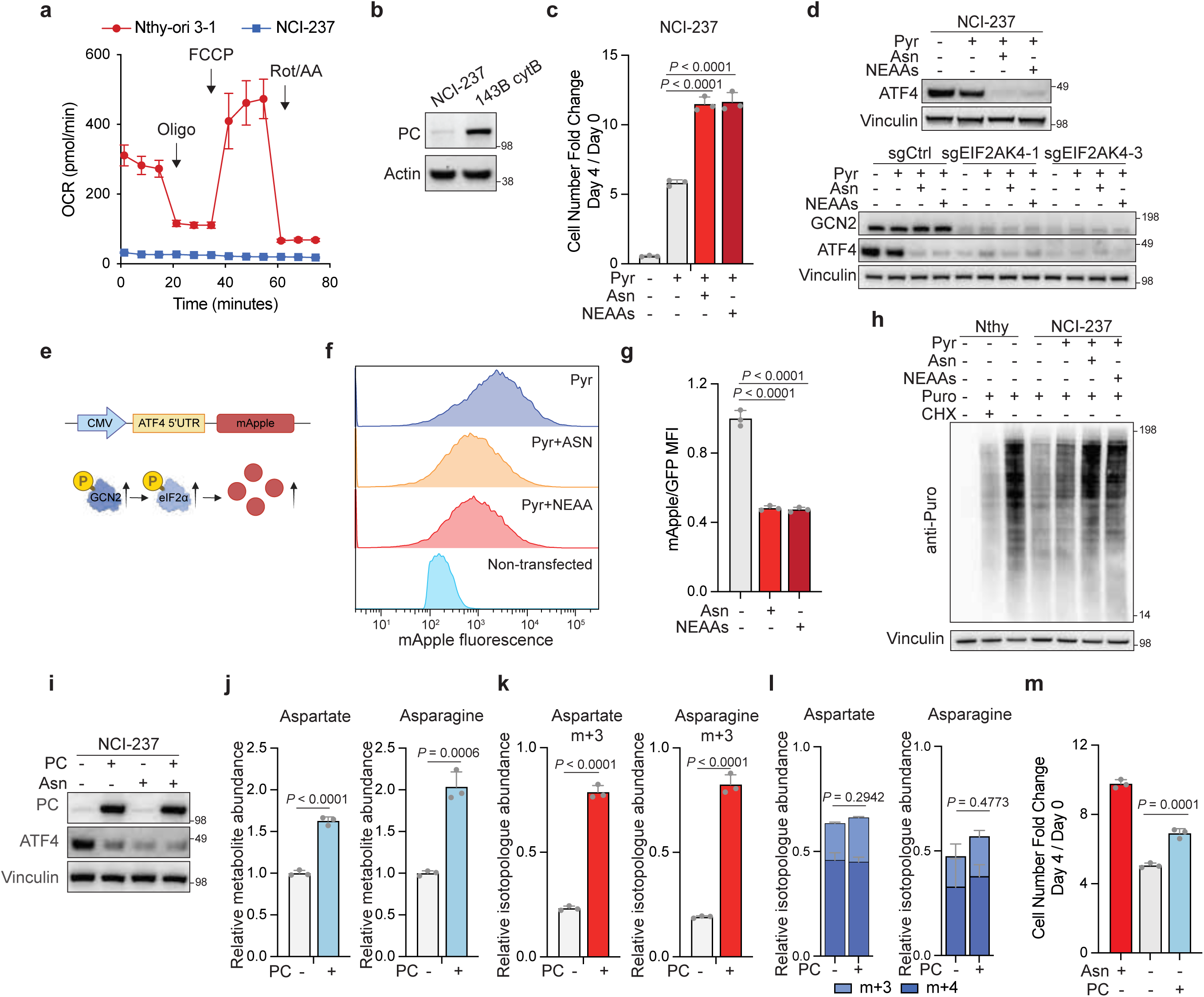
Patient-derived Hürthle cell carcinoma cells exhibit PC deficiency and asparagine auxotrophy. a, Oxygen consumption rate (OCR) of indicated cell lines measured by Seahorse Bioanalyzer. Oligo, oligomycin; Rot/AA, rotenone/antimycin A; FCCP, carbonyl cyanide-p-trifluoromethoxyphenylhydrazone. Values represent mean ± s.d. from n = 6 replicates. b, Immunoblot of PC expression levels in NCI-237 and 143B cytB cells. c, Cell proliferation measured as cell number fold change (day 4/day 0) in NCI-237 cells cultured in medium with 1 mM pyruvate, 100 µM asparagine or 100 µM NEAAs as indicated. d, Western blot of NCI-237 WT cells (top) and NCI-237 cells with sgCtrl (sgPRM1), sgEIF2AK4 (GCN2)-1 or sgEIF2AK4 (GCN2)-3 mediated CRISPR-Cas9 genome editing (bottom) cultured in medium with 1 mM pyruvate, 100 µM asparagine, or 100 µM NEAAs as indicated. e, Schematic illustrating the design of ATF4 reporter^55^. f, mApple fluorescence intensity distribution of NCI-237 reporter cells supplemented with 1 mM pyruvate and 100 µM asparagine or 100 µM NEAAs as indicated. mApple signal indicated translational activation of ATF4 based on the 5’UTR. The non-transfected group indicated cells without expression of ATF4 reporter. g, Normalized mApple/GFP median fluorescence intensity in NCI-237 reporter cells from (f), shown as the fold change relative to the group without asparagine or NEAAs supplementation. GFP was expressed as a transcriptional control. All three groups contained 1 mM pyruvate in the medium. h, Puromycin incorporation assay of NCI-237 and Nthy-ori 3-1 (Nthy) cells cultured in indicated medium for 24 h. Puromycin (90 µM) was added 10 min before sample collection, followed by immunoblotting with puromycin antibody. CHX, cycloheximide, a general translation inhibitor used at 10 µg/mL, was added 30 min before puromycin treatment as a control. i, Western blot of NCI-237 cells expressing empty vector control or PC cDNA, cultured with or without 100 µM asparagine supplementation as indicated. 1 mM pyruvate was present in all conditions. j, Relative total metabolite abundance of aspartate and asparagine in the NCI-237 cells expressing empty vector control or PC cDNA. 1 × 10^6^ cells were seeded for both conditions. After attachment, cells were cultured in glucose/glutamine-deficient DMEM containing 10 mM [U-^13^C] glucose + 1 mM [U-^13^C] pyruvate + 2 mM non-labeled glutamine for 24 h. Total metabolite abundance was calculated by summing the abundances of all isotopologue and normalized to the empty-vector control. k, Relative isotopologue abundance of aspartate (m+3) and asparagine (m+3) measured by LC-MS in the samples shown in j. Isotopologue abundance was calculated as the m+3 fractional enrichment × the corresponding relative total metabolite abundance shown in j. l, Relative isotopologue abundance of aspartate and asparagine (m+3 from reductive carboxylation or m+4 from oxidative TCA cycle) measured by LC-MS in NCI-237 cells expressing empty vector control or PC cDNA. 1 × 10^6^ cells were cultured in glucose/glutamine-deficient DMEM containing 2 mM [U-^13^C] glutamine + 10 mM non-labeled glucose + 1 mM non-labeled pyruvate for 24 h. The abundance of (m+3) and (m+4) aspartate and asparagine are presented from the same [U-^13^C] glutamine tracing experiment as indicated by the color scheme. m, Cell proliferation measured as cell number fold change (day 4/day 0) in NCI-237 cells, cultured in 1 mM pyruvate with 100 µM asparagine supplementation or PC expression as indicated. 200 µM uridine was supplemented in all the medium unless otherwise indicated (b-k). Data are shown as mean ± s.d. from n = 3 independent replicates unless otherwise indicated. Asn, asparagine. Pyr, pyruvate. Statistical significance was assessed using two-tailed t-tests (j, k, l), one-way ANOVA followed by Tukey’s multiple comparisons test (c, m) or Dunnett’s multiple comparisons test (g).

By contrast, breast cancer severity and metastasis potential has been correlated with high PC expression^52,53^. The highly aggressive triple-negative breast cancer (TNBC) cell line MDA-MB-231 exhibited high PC expression in comparison with RPE and 143B (Extended Data Fig. 7f). To test if other high-PC cancer cell lines, in addition to osteosarcoma-derived 143B cells, rely on endogenous PC-derived asparagine to proliferate, we deleted *PC* by CRISPR-Cas9 from MDA-MB-231 cells (Extended Data Fig. 7f). PC-deletion reduced the ability of MDA-MB-231 cells to proliferate *in vitro* even when grown in pyruvate- and uridine-supplemented medium and the reduced proliferation was significantly restored by asparagine addition in the PC-deleted cells. Asparagine supplementation exhibited no benefit in MDA-MB-231 WT cells under ETC inhibition (Extended Data Fig. 7g).

### Patient-derived Hürthle cell carcinoma cells exhibit PC deficiency and asparagine auxotrophy

The above results demonstrate that across multiple cell types and multiple mechanisms, both biochemical and genetic, the ability of respiratory-compromised cells to proliferate in the absence of exogenous asparagine is PC dependent. To further test whether these observations apply to patient-derived cell lines with mtDNA mutations under more physiologic conditions, we next studied NCI-237, a thyroid cancer patient-derived cell line that has been well characterized by others^50,51^. NCI-237 is a representative example of Hürthle cell carcinoma (HCC), a distinct subtype of thyroid cancer characterized by high heteroplasmic mitochondrial DNA mutations^23,24,54^. NCI-237 cells have a truncation point mutation in MT-ND5 at a heteroplasmy level of 90%^50,51^. To verify that this cell line had impaired respiration, we compared NCI-237 with the nontransformed thyroid cell line Nthy-ori 3-1, which is respiratory competent. In comparison to Nthy-ori 3-1 cells, NCI-237 exhibited low levels of basal respiration and showed little change in respiration rate when treated with ETC inhibitors or uncouplers (Fig. 7a). Additionally, NADH/NAD⁺ ratios were 4-fold higher in NCI-237 cells, consistent with impaired mitochondrial oxidation. As expected, the NADH/NAD⁺ ratios decreased upon supplementation with AKB or pyruvate (Extended Data Fig. 8a and 8b). NCI-237 cells, like RPE cells and Nthy-ori 3-1 cells, are epithelial and express a low level of PC protein relative to 143B cytB cells (Fig. 7b and Extended Data Fig. 7a). Like high heteroplasmic mtDNA^ΔScal^ and mtDNA^DdCBE-ND4^ RPE cells, either the NEAAs mixture or asparagine supplementation increased NCI-237 proliferation significantly in the presence of pyruvate and uridine (Fig. 7c). The other four amino acids (A, D, E, P) did not enhance proliferation (Extended Data Fig. 8c). NCI-237 cells exhibited high ATF4 expression in pyruvate- and uridine-supplemented medium. Adding asparagine or the mixture of NEAAs led to a significant reduction in ATF4 protein levels (Fig. 7d). GCN2-encoding gene *EIF2AK4* knockout by CRISPR-Cas9 also abolished ATF4 activation, regardless of pyruvate or asparagine supplementation, confirming that GCN2 is necessary for ATF4 activation (Fig.7d). Similar results were observed across different culture conditions that reduce reductive stress, including *Lb*NOX expression or AKB supplementation (Extended Data Fig. 8d-g).

**Fig. 8:**
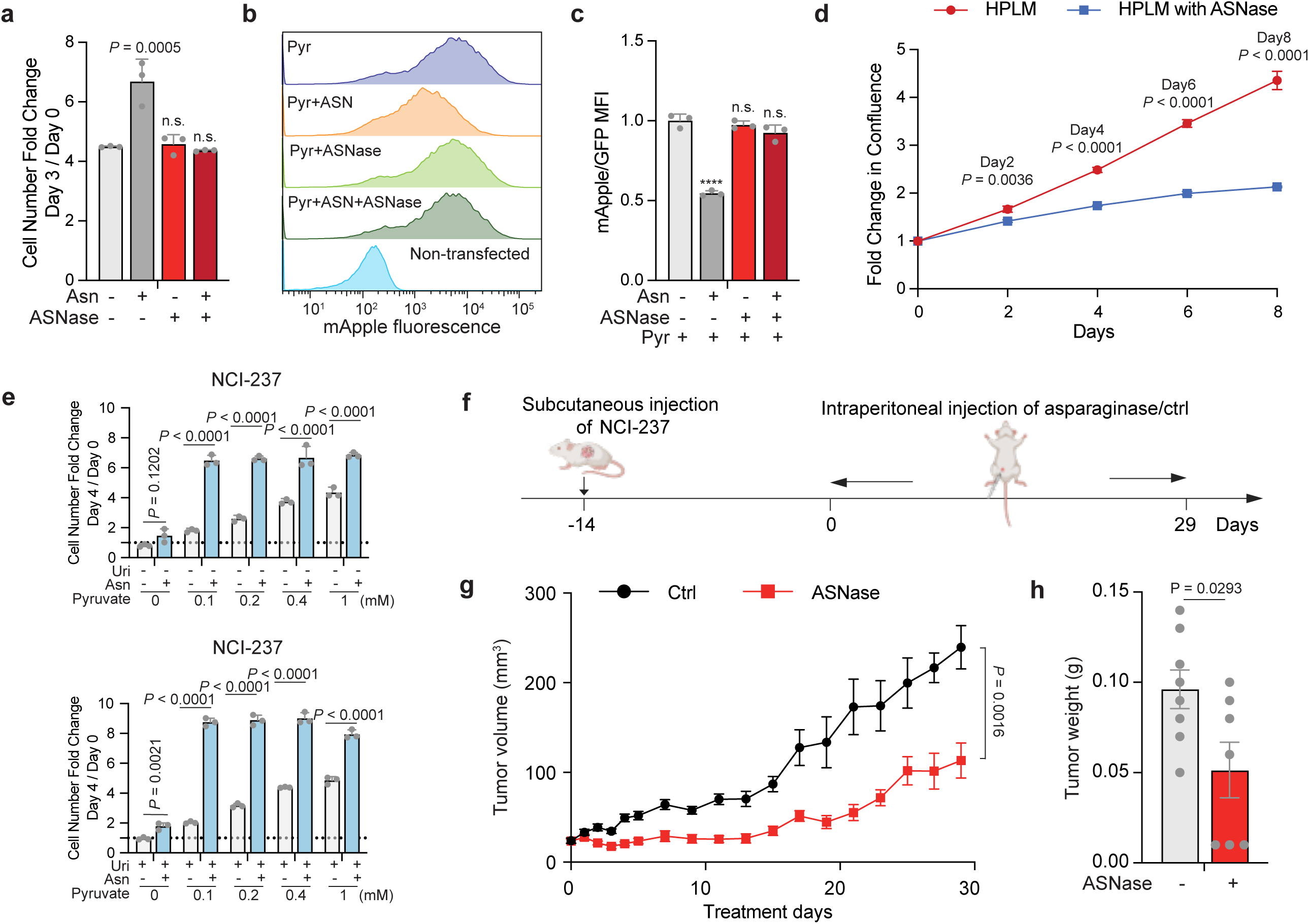
Asparaginase reduces Hürthle cell tumor growth *in vitro* and *in vivo.* a, Cell proliferation measured as cell number fold change of NCI-237 cells, cultured in 1 mM pyruvate with 100 µM asparagine supplementation or 0.1 IU/mL asparaginase as indicated. b, mApple fluorescence intensity distribution of NCI-237 reporter cells cultured in 1 mM pyruvate containing DMEM supplemented with 100 µM asparagine or 0.1 IU/mL asparaginase as indicated. The non-transfected group indicates cells without expression of ATF4 reporter. c, Normalized mApple/GFP median fluorescence intensity in NCI-237 reporter cells from (b), shown as the fold change relative to the group without asparagine or asparaginase treatment. d, Cell proliferation measured as fold change in confluence over the time of maintaining NCI-237 cells in HPLM supplemented with asparaginase 0.1 IU/mL as indicated. Medium was replaced daily. Statistical significance listed for fold change at each time point. e, Cell proliferation measured as cell number fold change of NCI-237 cells, cultured in DMEM without (top) or with (bottom) 200 µM uridine, supplemented with indicated concentrations of pyruvate, with or without 100 µM asparagine as indicated. f, Schematic illustrating the subcutaneous implantation of NCI-237 tumor xenograft and intraperitoneal injection of asparaginase or vehicle control. g, NCI-237 subcutaneous tumor xenograft growth curves (mm^3^) in mice treated with asparaginase (80 IU/mouse/day, n = 7) or vehicle control (n = 8) from treatment start date through endpoint. Tumor volumes were measured by caliper measurements. The data are representative of 2 independent experiments. h, Endpoint tumor weight (g) of NCI-237 tumor xenograft from (f). 200 µM uridine was supplemented in a-c. Data are shown as mean ± s.d. from n = 3 independent replicates in a, c, d, e, and as mean ± s.e.m. in g-h. Statistical significance was assessed using two-tailed t-tests (d, g, h), one-way ANOVA followed by Dunnett’s multiple comparisons test (a, c), or two-way ANOVA followed by Šidák multiple comparisons test (e). ASNase, asparaginase.

To test whether asparagine suppression of ATF4 involved a change in ISR-regulated translation, we utilized a previously published ATF4 reporter^55^. ISR upstream kinases phosphorylate eIF2α, which selectively translates mRNAs containing ISR-responsive elements in their 5’ untranslated region (5’UTR). The reporter employs the 5’UTR of ATF4, followed by the mApple coding sequence in place of the ATF4 coding sequence and acts as a reporter for ISR activation levels (Fig. 7e). NCI-237 cells expressing the ATF4 reporter exhibited significant mApple fluorescence when cultured in DMEM supplemented with uridine and pyruvate, which was reduced by either asparagine or NEAAs supplementation (Fig. 7f). The ratio of median fluorescence intensity of mApple/GFP decreased significantly with asparagine or NEAAs supplementation (Fig. 7g). Either asparagine or NEAAs supplementation also enhanced translation in NCI-237 cells as measured by the puromycin incorporation assay (Fig. 7h).

To test whether PC expression could bypass the need for exogenous asparagine, we ectopically expressed PC in NCI-237 cells (Fig. 7i-m). PC expression led to reduced ATF4 protein levels (Fig. 7i). PC-expressing NCI-237 cells exhibited increased levels of aspartate (1.6 ± 0.1)-fold and asparagine (2.0 ± 0.2)-fold (Fig. 7j). In PC-expressing cells, the relative contribution of aspartate (m+3) derived from [U-^13^C]-glucose + [U-^13^C]-pyruvate labeling increased 3.4-fold and asparagine (m+3) increased 4.3-fold in comparison to NCI-237 control cells (Fig. 7k). The labeling percentage of PC-derived aspartate (m+3) and asparagine (m+3) as well as TCA metabolites citrate (both m+5 and m+3), malate (m+3), and fumarate (m+3) were also all significantly increased in the PC-expressing NCI-237 cells (Extended Data Fig. 8h).

In NCI-237 cells, which carry a high heteroplasmic mtDNA mutation in Complex I, [U-^13^C] glutamine contributed significantly to de novo aspartate and asparagine production through both oxidation in the TCA cycle (m+4) and reductive carboxylation (m+3). The total amount of both aspartate and asparagine labeled from glutamine also increased upon PC expression, but the difference is not statistically significant (Fig. 7l). Together, with the above experiments, this suggests that de novo synthesis from glucose + pyruvate and de novo synthesis from glutamine combined contribute up to 89% of the free aspartate pool and 80% of the free asparagine pool (Fig. 7j-l and Extended Data Fig. 8h). Along with elevated cellular levels of asparagine and aspartate, PC ectopic expression significantly promoted cell growth (Fig. 7m).

### Asparaginase reduces Hürthle cell tumor growth *in vitro* and *in vivo*

Extracellular asparagine can be depleted by the addition of recombinant L-asparaginase which has been used in the treatment of acute lymphoblastic leukemia for decades^56–59^. To evaluate whether NCI-237 cells depend on extracellular asparagine, we tested the effect of L-asparaginase on cell proliferation (Fig. 8). In vitro, treatment with L-asparaginase completely abolished the proliferative effect of extracellular asparagine (Fig. 8a). As measured by the ATF4 reporter assay described above, treatment with extracellular L-asparaginase also reversed the ability of asparagine to suppress ATF4 translation as measured by mApple fluorescence (Fig. 8b and 8c).

A physiologic medium mimicking the concentrations of polar metabolites in human plasma was used to assess the effect of asparagine depletion by L-asparaginase on NCI-237 cell proliferation. L-asparaginase significantly reduced proliferation of NCI-237 cells cultured in Human Plasma Like Medium (HPLM) (Fig. 8d). The difference in cell proliferation was statistically significant by Day 2 and at every subsequent time point measured (Fig. 8d). The 8-day time scale is presented to demonstrate that cells do not exhibit evidence of compensatory adaptation to restore cell proliferation when depleted of exogenous asparagine. In the HPLM experiments, the medium was replaced daily to avoid unintended depletion of other required nutrients. As L-asparaginase can also potentially deamidate glutamine, we also tested the effect of addition of 0.1 IU/mL L-asparaginase on glutamine and asparagine levels in the HPLM medium. The HPLM medium ± 0.1 IU/mL L-asparaginase was assayed after 24 hours at 37 °C. No statistically significant difference in glutamine levels was detected between L-asparaginase-treated medium and control medium. In contrast, asparagine was readily detected in control HPLM but was not detected in L-asparaginase treated medium (Extended Data Fig. 8i).

The ability of NCI-237 cells to grow in HPLM medium was surprising as this medium is not supplemented with supraphysiologic levels of pyruvate and uridine that are normally utilized for maintaining respiratory-compromised cells *in vitro*. However, the medium does contain a physiologic level of pyruvate. Our initial experiments had identified the ability of asparagine to enhance the cell proliferation in medium supplemented with exogenous pyruvate and uridine. The HPLM experiment result prompted us to revisit the requirements for pyruvate and uridine further. A pyruvate titration experiment in NCI-237 cells demonstrated that the proliferative dependence on physiologic asparagine levels was even more pronounced at physiologic pyruvate levels (0.1 mM) than at supraphysiologic pyruvate levels. Asparagine addition increased cell proliferation at a physiologic level (0.1 mM) of pyruvate by more than four-fold while the presence of exogenous asparagine increased proliferation less than two-fold in medium containing a supraphysiologic level (1 mM) of pyruvate. Asparagine alone had no significant effect on cell proliferation in the complete absence of exogenous pyruvate. However, once cells are provided with physiologic levels of pyruvate, physiologic levels of asparagine are sufficient to maximize cell proliferation. Addition of uridine enhanced overall proliferation but does not influence the relative abilities of pyruvate and asparagine to promote cell proliferation across the range of pyruvate levels tested (Fig. 8e).

NCI-237 cells form robust xenograft tumors in immunodeficient mice where they are not exposed to supraphysiologic levels of pyruvate and uridine^50,51^. To assess whether the *in vivo* growth of NCI-237 was dependent on extracellular asparagine, we established xenografts in immunodeficient NSG mice. Two weeks after subcutaneous injection of NCI-237 cells, mice with established tumors were randomized to receive either L-asparaginase (80 IU/mouse) or a vehicle control daily for 4 weeks. This dose was chosen because pharmacokinetic experiments in mice have demonstrated that this dose depletes extracellular asparagine without significantly affecting systematic glutamine^60,61^. Tumor volume was monitored every other day. Following the treatment period, animals were sacrificed, and the tumors resected and weighed (Fig. 8f). Asparaginase treatment significantly reduced tumor volumes and end-point tumor weights (Fig. 8g and 8h), indicating that NCI-237 tumor growth depends on extracellular asparagine availability.

## Discussion

The present findings demonstrate that increasing levels of mtDNA^ΔScal^ heteroplasmy lead to a progressive decline in both basal oxygen consumption and maximal respiratory capacity. At heteroplasmy exceeding 85%, cells exhibit a threshold-dependent induction of the ISR, which can be rescued by supplementation with exogenous asparagine. This heteroplasmy-dependent asparagine auxotrophy is observed in both large-scale mtDNA deletion models and in isogenic point mutation models affecting a single Complex I subunit (ND4), indicating that the asparagine auxotrophy is not restricted to multi-gene disruptions. Consistent with the mtDNA heteroplasmy results, respiratory deficiency induced by pharmacologic ETC inhibition or genetic ablation of nuclear-encoded ETC components also results in dependence on extracellular asparagine, supporting that impaired TCA cycle-coupled electron transport chain activity, rather than specific mtDNA lesions, causes the asparagine deficiency. Ectopic expression of PC overcame asparagine auxotrophy in respiratory-deficient cells, including RPE cells with high heteroplasmic mtDNA^ΔScal^ deletion or mtDNA^ddcbe-ND4^ mutation, as well as patient-derived NCI-237 cells with high heteroplasmic Complex I mutation, and ETC-inhibited thyroid and colorectal cancer cell lines lacking mtDNA mutations. In contrast, in 143B-derived cybrids or breast cancer cell line MDA-MB-231, which constitutively express PC, genetic deletion of PC reduced asparagine synthesis and activated ISR. These results demonstrate that mitochondrial PC activity promotes asparagine generation in cells with mtDNA mutations at high heteroplasmy.

Asparagine serves a unique role in murine and human cells, as it is exclusively used to charge tRNA^Asn^ for protein synthesis but cannot be metabolized into other metabolic intermediates, including aspartate^34^. The presence of any uncharged tRNA is sufficient to activate GCN2 kinase activity. GCN2-dependent phosphorylation of eIF2α reduces overall translation while selectively enhancing ATF4 expression^37,62^. ATF4 is a transcription factor that induces stress-responsive genes, including asparagine synthetase (ASNS)^63–65^. Our proteomic analysis highlights ASNS as a prominent ATF4 target (Fig. 3a). Despite the high level of ATF4-dependent ASNS protein expression, cells with high heteroplasmic mtDNA mutations are unable to produce sufficient asparagine to support the synthesis of proteins needed for cell growth and proliferation. The high demand for aspartate in the synthesis of nucleotides may prevent the accumulation of sufficient aspartate in the cytosol to support *de novo* asparagine synthesis in sufficient quantities to meet cellular demand in respiratory-deficient cells. While cellular metabolism has long been considered to follow the law of mass action with the whole cell represented as a homogeneous environment, it is increasingly clear that different intracellular compartments and organelles contain distinct levels of metabolites. For example, recent studies have demonstrated that mitochondrial aspartate can be channeled into nucleotide production^4^, such a process would compromise the ability of respiratory-deficient cells to produce enough cytosolic aspartate to support *de novo* asparagine synthesis. When respiratory compromise limits glutamine anaplerosis-dependent support of aspartate/asparagine production, mitochondrial PC-dependent conversion of pyruvate to oxaloacetate appears capable of compensating. Mitochondrial respiration deficiency-induced ISR has been described in various contexts. Prior studies have shown that in proliferating cells, including myoblasts, ETC inhibition activates the ISR predominantly through GCN2 in response to asparagine insufficiency, whereas in differentiated, non-proliferating cells, ISR activation is linked to energetic stress^66^. Consistent with these prior findings, our results suggest that proliferating cells, such as cancer cells, are sensitive to asparagine deficiency due to a sustained demand for net protein synthesis. In contrast, non-proliferating cells have a reduced anabolic demand and can recycle intracellular asparagine through protein turnover^67^.

The present findings underscore the importance of oxidative phosphorylation in enabling glutamine anaplerosis to support sufficient aspartate production for both nucleic acid and amino acids syntheses^4^. Previous studies have suggested that asparagine synthesis is sensitive to intracellular aspartate production^34,68^. This is in part due to the fact that aspartate is poorly taken up from the extracellular environment and is present at 50-100 µM in extracellular fluid^30–32^. Past studies have demonstrated that supraphysiologic levels of extracellular aspartate are required to

restore the intracellular aspartate-dependent syntheses needed to support the proliferation of respiration-compromised cells in the absence of pyruvate. We have confirmed that finding. However, we found that cells with high heteroplasmy loss-of-function mtDNA mutations can synthesize sufficient aspartate to maintain other aspartate-dependent processes required for cell growth when redox imbalance is corrected. As shown in Figure 8, when cultured in medium containing physiologic levels of asparagine and pyruvate, the cell’s dependence on extracellular asparagine could be exploited to suppress the growth of respiratory-deficient cancer cells by L-asparaginase treatment both *in vitro* and *in vivo*.

Most cancer cells lack mitochondrial DNA mutations that compromise respiration^25^. Several common oncogenic mutations including activating mutations in PI3K, AKT, and Myc enhance glutamine uptake and metabolism^69,70^. In such transformed cells, mitochondrial metabolism of glutamine in the TCA cycle supports *de novo* aspartate and asparagine synthesis, a process that requires electron transport chain activity. Based on this, it has recently been suggested that combining ETC inhibitors with L-asparaginase may prevent TCA cycle-dependent conversion of glutamine into asparagine in respiration-competent cells^71^. Unfortunately, currently available ETC inhibitors that are safe for systemic administration are generally not effective enough at suppressing TCA cycle-coupled OXPHOS to levels that impair asparagine synthesis. However, there are an increasing number of tumors with high heteroplasmic mutations in mtDNA being identified. Pan-cancer analyses have demonstrated that a subset of mutations, particularly truncating variants in OXPHOS genes, can expand to high and even near-homoplasmic levels. Such high-heteroplasmy mutations are enriched in specific tumor types, including thyroid, renal, and colorectal cancers^23–25,49^. Hürthle cell carcinomas are characterized by recurrent mtDNA mutations that frequently reach near-homoplasmic levels, indicating that heteroplasmy levels exceeding 85% are not uncommon in this tumor type^72^. The present data suggest that L-asparaginase may have efficacy in this subset of tumors.

Most proliferating cells do not express high levels of pyruvate carboxylase. Pyruvate carboxylase is normally expressed in differentiated cells utilizing glucose rather than glutamine for TCA cycle anaplerosis^73^. NCI-237, a patient-derived Hürthle cell carcinoma line with high mtDNA heteroplasmy lacks significant PC activity and exhibits asparagine auxotrophy. L-asparaginase, which converts extracellular asparagine to aspartate, reduced NCI-237 tumor cell growth both *in vitro* and *in vivo*, confirming that the tumor cells depend on extracellular asparagine. L-asparaginase has proven to be an effective part of curative therapy of acute lymphoblastic leukemia (ALL), but its use in other tumors has been limited by the immunogenicity of L-asparaginase and the side effects exhibited particularly when combined with other drugs. With the exception of ALL, other cancer types have not previously been shown to be auxotrophic for asparagine^57,59^. From the findings reported here, tumors with high heteroplasmic mtDNA mutations and low PC expression are auxotrophic for asparagine. These mtDNA-mutant tumors are potential new targets for L-asparaginase therapy, warranting further optimization of L-asparaginase as a therapeutic.

## Methods

### Cell lines and cell culture

The ARPE-19 (RPE) wild-type and RPE mtDNA^ΔScal^ isogenic cell lines were obtained from Dr. Agnel Sfeir^29^. The 143B cytB cells were a gift from Dr. Ralph J. DeBerardinis at UT Southwestern and were previously established by Rana et al.^28^. The 143B UQCRC2 KO and COX4I1 KO cells were a gift from Dr. Jessica B. Spinelli at UMass Chan Medical School^74^. NCI-237 cells and TPC1 cells were a gift from Dr. David McFadden at UT Southwestern^38^. The 143B, DLD1, MDA-MB-231 wild-type cells were obtained from American Type Culture Collection (ATCC). Nthy-ori 3-1 cells were obtained from Millipore (#90011609).

All cell lines were maintained in Dulbecco’s Modified Eagle Medium (DMEM) supplemented with 10% fetal bovine serum (FBS), 200 µM uridine, 100 units/mL penicillin, 100 µg/mL streptomycin, 1 mM sodium pyruvate, and 100 µM non-essential amino acids (NEAAs), as previously reported^29^. This medium is referred to as complete medium and was used for cell maintenance and for initial seeding in all experiments unless otherwise specified. All cultures were incubated at 37°C in an atmosphere containing 20% oxygen and 5% CO₂, and were regularly confirmed to be mycoplasma-free using the MycoAlert Mycoplasma Detection Kit (Lonza, LT07-318).

### Mice

The mice NOD.Cg-Prkdc^scid^ Il2rg^tm1Wjl^/SzJ strain #005557 were purchased from Jackson Laboratories and bred in-house. All animal experiments described adhered to policies and practices approved by Memorial Sloan Kettering Cancer Center’s Institutional Animal Care and Use Committee (IACUC) and were conducted in accordance with NIH guidelines for animal welfare (Protocol Number 11-03-007, Animal Welfare Assurance Number FW00004998).

### Plasmids

#### Generation of knockout constructs

Gene knockout via CRISPR-Cas9 was carried out using the lentiCRISPR v2 system (Addgene, 52961 and 98291). The human control sgRNA (sgCtrl) targets the non-expressed, silent gene *PRM1* and was used to induce genome cutting without affecting active genes. The gRNA sequences are in Supplementary Table 4.

#### Generation of asparagine reporter constructs

The polyASN-BFP construct was generated using the doxycycline-inducible lentiviral vector pInducer20 (Addgene, 44012). The vector was digested with XhoI and NheI, followed by gel purification. A synthesized gene fragment containing a start codon (ATG), a poly-asparagine tract encoding eight asparagines, a start codon-less BFP preceded by a flexible linker (GGGGSGGGGS), a degron sequence, and a 27 bp overlap with the pInducer20-neomycin region was obtained from Twist Bioscience. The synthesized fragment was PCR-amplified and inserted into pInducer20-neo using Gibson assembly. The sequence of the synthesized fragment is in Supplementary Table 4.

#### Generation of PC gRNA resistant construct

A point mutation was introduced into the pBABE-neomycin-PC-MYC construct (Addgene, 184550) via site-directed mutagenesis to prevent guide RNA targeting while maintaining the wild-type protein sequence. The mutation was performed using PfuTurbo DNA polymerase (Agilent, 600250). The primers used for generating the point mutation are in Supplementary Table 4.

#### Generation of LbNOX inducible expression construct

cDNA for Flag-tagged LbNOX was obtained from Addgene (75285) and cloned into the pINDUCER20 (Addgene, 44012) tet-on lentiviral expression system.

#### Generation of stable cell lines

Lentiviral particles were generated in HEK293T cells by co-transfecting the viral vector with psPAX2 and pCMV-VSV-G packaging plasmids (Addgene). Retroviral particles were generated with the packaging plasmids pCG-gag-pol and pCMV-VSV-G (Addgene). After collection, viral supernatants were filtered through 0.45-μm nylon membranes and used to transduce target cells in the presence of 8 μg/mL polybrene (Sigma). At 24 h post-transduction, cells were subjected to antibiotic selection with either puromycin (2 μg/mL, Sigma), hygromycin (200 μg/mL, InvivoGen), or neomycin (1 mg/mL, InvivoGen), depending on the construct. Antibiotic-resistant populations were established before further experiments. For 143B cytB PC knockout cells, single cells were sorted into individual wells to derive clonal populations, which were then expanded and screened for successful PC knockout.

### mtDNA deletion quantified by droplet digital PCR (ddPCR)

Extraction of genomic DNA:

Total genomic DNA was purified using the DNeasy Blood & Tissue Kit (Qiagen, 69504) following the manufacturer’s protocol for mtDNA heteroplasmy measurements. Cell pellets were resuspended in 200 μL PBS, followed by the addition of 20 μL Proteinase K and 200 μL Buffer AL. The mixture was vortexed and incubated at 56 °C for 10 min. DNA was then purified using spin columns. Genomic DNA samples were normalized to equal concentrations and subjected to ddPCR.

Assay design for ddPCR:

Assays specific for the detection of MT-CO3, ACTB, and a control region of the mitochondrial genome (ddMDM, within the D-loop region) were designed and ordered through Bio-Rad. Cycling conditions were tested to ensure optimal annealing/extension temperature as well as optimal separation of positive from empty droplets. Optimization was done with a known positive control. After PicoGreen quantification, 0.1 ng DNA were combined with locus-specific primers, FAM- and HEX-labeled probes, HindIII, and digital PCR Supermix for probes (no dUTP). All reactions were performed on a QX200 ddPCR system (Bio-Rad) and each sample was evaluated in technical duplicates or triplicates. Reactions were partitioned into a mean of ∼20,000 droplets per well using the QX200 droplet generator. Emulsified PCRs were run on a 96-well thermal cycler using cycling conditions identified during the optimization step (95 °C for 10 min; 40 cycles of 94 °C for 30 s and 56 °C for 1 min; 98 °C for 10 min; hold at 4 °C). Plates were read and analyzed with the QuantaSoft software to assess the number of droplets positive for gene of interest, reference gene, both, or neither.

### Generation of mtDNA^DdCBE-ND^^4^ heteroplasmic cell lines

The plasmids ND4.2-Right DdCBE-G1397-C-T1413I-GFP (Addgene, 179686) and ND4.2-Left TALE-G1397-N-DddA11-mCherry (Addgene, 179682) were a gift from Dr. David R. Liu and have been described previously^42,43^. Plasmids were transfected into RPE wild-type cells using a 4D-Nucleofector X unit (Lonza) with SF Nucleofection Buffer and the ER-100 program. Three days after transfection, GFP- and mCherry-double-positive cells were single-cell sorted to derive clonal populations with defined levels of heteroplasmy and maintained in DMEM supplemented with pyruvate, uridine, and NEAAs. mtDNA heteroplasmy at the targeted point mutation was quantified by extracting genomic DNA as described above, followed by amplicon sequencing. Briefly, primers were designed to amplify the mtDNA region containing the mutation, and PCR was performed using standard cycling conditions (see Supplementary Table 4 for primer sequences). PCR products were gel-purified and subjected to Sanger sequencing to determine heteroplasmy levels in mtDNA^DdCBE-ND4^ cell lines.

### Western blot

Cells were seeded in complete medium and cultured for 24 h to let cells attach. 24 h after seeding, cells were changed to experimental medium as indicated. For harvest, cells were washed twice with ice-cold phosphate-buffered saline (PBS) and lysed using RIPA buffer containing protease and phosphatase inhibitors. Lysates were incubated at 4°C for 30 minutes to facilitate protein extraction, then clarified by centrifugation. Protein concentration was determined by Bradford protein assay (Bio-Rad, 5000006), following which equal amounts of protein were loaded and separated by SDS–polyacrylamide gel electrophoresis (SDS–PAGE). Proteins were transferred to nitrocellulose membranes, except that phospho-GCN2 (T899) blot was transferred to PVDF membranes. Membranes were blocked with 5% non-fat milk in TBST and incubated with primary antibodies in 3% bovine serum albumin (BSA) in TBST. Afterward, they were incubated with anti-rabbit HRP-conjugated IgG (1:4,000) or anti-mouse HRP-conjugated IgG (1:3,000). Membranes were washed with TBST between incubations. Signals were detected using an ECL detection reagent (Thermo Fisher Scientific, PI32106) and a ChemiDoc MP Imaging System (Bio-Rad). Western blot images were analyzed using Image Lab (v.6.1). All antibodies used for Western blotting are listed in Supplementary Table 2.

### Proliferation assay

Cell number-based fold change: Cell proliferation experiments shown by cell number fold change were conducted by seeding cells in the complete medium used for cell maintenance. Cells were plated at a density of 7.5 × 10^3^ to 15 × 10^3^ cells per well in 12-well plates. The following day, cells were washed once with warm PBS, and the medium was replaced with the experimental medium consisting of DMEM supplemented with 10% dialyzed FBS (dFBS, Gemini), 100 units/mL penicillin, 100 µg/mL streptomycin, and 200 µM uridine. Where indicated, 1 mM sodium pyruvate, 100 µM indicated amino acids were added. Cell number counts at the start and end of the experiments were measured in triplicate wells using a Multisizer 3 Coulter Counter (Beckman Coulter).

Confluence-based fold change: Cells were plated at a density of 5 × 10^3^ cells per well in 24-well plates. After replacement with experimental medium as described above in cell number-based fold change, images were acquired at time 0 and at the indicated time points on the Incucyte S3 Live-cell analysis system (Sartorius) with a 10x objective in brightfield mode. 36 fields per well were captured at each time point. Cell confluence was measured using Incucyte analysis software with settings as follows: 0.8 segmentation adjustment, 400 μm^2^ hole fill, 700 μm^2^ minimal area.

Proliferation assay with HPLM (Fig. 8d): NCI-237 cells were adapted in HPLM medium supplemented with 10% FBS, 100 units/mL penicillin, and 100 µg/mL streptomycin. 2 × 10^4^ cells were seeded per well in 6-well plates. The following day, cells were washed once with warm PBS, and the medium was replaced with HPLM supplemented with 10% dFBS, 100 units/mL penicillin, 100 µg/mL streptomycin. For the asparaginase treated conditions, 0.1 IU/mL asparaginase was added to the medium. Both control and asparaginase supplemented media were replaced daily throughout the experiments. Cell proliferation was monitored every other day by Incucyte S3 Live-cell analysis system as described above.

### Flow cytometry

For asparagine-reporter and ATF4-reporter measurements, cells were seeded at 3 × 10^4^ cells per well in 12-well plates in the complete medium used for cell maintenance. The following day, cells were washed once with warm PBS, and the medium was replaced with experimental conditions. For asparagine-reporter measurement, all treatment media contained 200 ng/mL doxycycline to induce reporter expression. After 24 h, cells were harvested with trypsin, washed twice with PBS, and stained with Zombie NIR^TM^ dye for 15 min at room temperature. Flow cytometry was performed using a BD LSRFortessa, and data were analyzed with FlowJo.

### Oxygen consumption measurements

Oxygen consumption rate (OCR) was measured using the XFe96 Extracellular Flux Analyzer (Agilent) following the manufacturer’s protocol. Cells were seeded at a density of 3 × 10^4^ cells per well in Seahorse microplates (Agilent) in the complete medium used for cell maintenance and allowed to adhere overnight. After removing the culture medium, cells were incubated with Seahorse XF DMEM medium (Agilent, 103575) supplemented with 10 mM glucose, 2 mM glutamine, and 1 mM pyruvate. OCR measurements were taken under basal conditions, followed by sequential injections of oligomycin (1 μM), FCCP (0.5 μM), and a rotenone/antimycin A mix (1 μM each) as indicated. OCR values were normalized to cell numbers as counted by Multisizer 3 Coulter Counter (Beckman Coulter) after the Seahorse assay.

### NADH/NAD^+^ measurements using enzymatic assay

The NADH/NAD^+^ ratio was quantified using the NAD^+^/NADH-Glo Assay kit (Promega, G9071), following the manufacturer’s instructions. Cells (3 × 10^4^) were seeded in six-well plates in the complete medium used for cell maintenance. The next day, cells were washed with PBS, and the medium was replaced with experimental medium (DMEM supplemented with 10% dialyzed FBS, 100 units/mL penicillin, 100 µg/mL streptomycin, and 200 µM uridine) along with additional metabolites as specified. After 24 hours, cells were washed twice with ice-cold PBS and collected by adding 1 mL of ice-cold 80% methanol over dry ice. The samples were incubated overnight at -80°C, then centrifuged at 21,000g for 30 minutes at 4°C. The supernatants were dried in a vacuum evaporator (Genevac EZ-2 Elite) for 6 hours. The dried metabolites were resuspended in buffer A (0.2 N NaOH diluted 1:1 with PBS) as the lysate. For NAD^+^ measurement, 20 μL of the lysate was transferred to a PCR tube, mixed with 20 μL buffer A and 20 μL of 0.4 N HCl, and incubated at 60°C for 30 minutes. For NADH measurement, 20 μL of the lysate was added to PCR tubes and incubated at 75°C for 1 hour. The acidic conditions selectively degrade NADH, and the basic conditions degrade NAD^+^. After incubation, samples were neutralized with 20 μL of neutralizing solution, which is 0.5 M Tris base for NAD^+^ or 0.25 M Tris in 0.2 N HCl for NADH, and mixed 1:1 with NAD^+^/NADH-Glo detection reagent. After 30 minutes of incubation at room temperature, luminescence was detected using a plate reader.

### Metabolite analysis using GC-MS

Cells were seeded in six-well plates 48 hours prior to collection and washed once with PBS before changing the medium. To assess metabolite levels under conditions with or without 1 mM sodium pyruvate, the medium was replaced 24 hours post-seeding, and cells were cultured in the respective medium for the indicated time points. Metabolism was quenched by adding 1 mL of ice-cold 80% methanol containing 2 µM deuterated 2-hydroxyglutarate (d5-2HG) and storing samples at -80°C overnight. The methanol-extracted metabolites were collected and centrifuged at 21,000g for 30 minutes to remove protein debris. The supernatant was evaporated using a vacuum concentrator (Genevac EZ-2 Elite) for 6 hours. The dried metabolites were resuspended in 20 mg/mL methoxyamine hydrochloride (Sigma, 226904) in pyridine (Thermo Fisher Scientific, TS-27530) and incubated for 90 minutes at 30°C. They were then derivatized using MSTFA containing 1% TMCS (Thermo Fisher Scientific, TS-48915) for 30 minutes at 37°C. Metabolite analysis was performed on an Agilent 7890A GC coupled with an Agilent 5975C Mass Selective Detector using electron impact ionization. The GC was operated in splitless mode with a constant helium flow of 1 mL per minute. One microliter of derivatized sample was injected onto an HP-5MS column (15 m × 0.25 mm, 0.25 µm film thickness), with an inlet temperature of 250°C and a temperature gradient from 60°C to 290°C over 25 minutes. Data were processed using Mass Hunter Quantitative Analysis software (v.10.0, Agilent Technologies). Metabolite peak areas were normalized to the internal standard (d5-2HG) and biomass (cell count × average cell size as determined by Coulter counter measurements). Ions used for quantifying steady-state metabolites included: d5-2HG (m/z 354), aspartate (m/z 232), asparagine (m/z 116), citrate (m/z 273), and malate (m/z 233). All chromatographic peaks were manually verified by comparison to reference spectra for each metabolite.

For the measurement of glutamine and asparagine in HPLM following asparaginase treatment, HPLM supplemented with 10% dialyzed FBS (dFBS), 100 units/mL penicillin, and 100 µg/mL streptomycin was prepared with or without 0.1 IU/mL asparaginase, matching the culture conditions used in Fig. 8d. 2 mL of medium was added to each well of cell-free 6-well plates, which were maintained in the same cell-culture incubator for 24 hours. Following incubation, 200 µL of medium from each well was collected and mixed with 1 mL of ice-cold methanol. Samples were centrifuged to precipitate proteins and remove debris derived from dFBS, and the resulting metabolite-containing supernatants were collected. Metabolite extraction and mass-spectrometric analysis were then performed as described above. Ions used for relative metabolites abundance included: m/z 347 for glutamine and m/z 116 for asparagine. Peak identities were confirmed using qualifier ions at m/z 156 and 245 for glutamine and m/z 231 and 333 for asparagine. Metabolite standards were analyzed in parallel to confirm peak identity. All chromatographic peaks were manually verified by comparison to reference spectra for each metabolite.

### Metabolite analysis using LC-MS

Cells were seeded in six-well plates 40 hours prior to the tracing experiment and washed once with PBS before changing the medium. For [U-^13^C] glucose tracing, the experimental medium was made from glucose-deficient DMEM supplemented with 10 mM [U-^13^C] glucose, 1 mM AKB and 200 μM uridine. For [U-^13^C] glutamine tracing, the experimental medium was made from glutamine-deficient DMEM supplemented with 2 mM [U-^13^C] glutamine, 1 mM pyruvate and 200 μM uridine. For [U-^13^C] glucose and [U-^13^C] pyruvate tracing, the experimental medium was made from glucose-deficient DMEM supplemented with 10 mM [U-^13^C] glucose, 1 mM [U-^13^C] pyruvate and 200 μM uridine. Tracer incubation times were 6 h for [U-^13^C] glucose tracing, 8 h for [U-^13^C] glutamine tracing, and 24 h for the indicated [U-^13^C] glucose/[U-^13^C] pyruvate and [U-^13^C] glutamine tracing experiments in RPE cells or NCI-237 cells. Exact medium composition and durations are specified in the corresponding figure legends. Metabolism was quenched and metabolites were extracted by removing the medium and adding 1 mL of 80% methanol chilled to -80 °C. After overnight incubation at -80 °C, cells were collected and centrifuged at 21,000g for 30 min at 4 °C. Supernatants were collected, then dried down and re-dissolved in water. Targeted LC/MS analyses were performed on a Q Exactive Orbitrap mass spectrometer (Thermo Scientific) coupled to a Vanquish UPLC system (Thermo Scientific). The Q Exactive operated in polarity-switching mode. For certain samples, only negative mode was used with a selected ion monitoring scan to improve the detection of asparagine. A Sequant ZIC-pHILIC column (2.1 mm i.d. × 150 mm, particle size of 5 µm, Millipore Sigma) was used for separation of metabolites. A 2.1 × 20 mm guard column with the same packing material was used for protection of the analytical column. Flow rate was set at 150 μL/min. Buffers consisted of 100% acetonitrile for mobile phase A, and 0.1% NH_4_OH/20 mM CH_3_COONH_4_ in water for mobile phase B. The chromatographic gradient ran from 85% to 30% A in 20 min followed by a wash with 30% A and re-equilibration at 85% A. The raw data was processed using El-MAVEN (v0.12.0). Metabolites and their isotopologues were identified on the basis of exact mass within 5 ppm and standard retention times.

### Proteomic assay

Cells were seeded in 15 cm plates 24 h prior to treatment with experimental conditions. On the next day of seeding, cells were washed with warm PBS once and changed into experimental medium, DMEM supplemented with 10% dFBS, 200 µM uridine, and 1 mM pyruvate. 100 µM asparagine was added in experimental medium as indicated. Cells were cultured in the respective media for 48 hours before harvesting for cell pellets.

### Protein digestion

The cell pellets were resuspended in 50 μL of 8 M Urea and 50 mM EPPS (pH 8.5), then treated with tris(2-carboxyethyl)phosphine (TCEP) at a final concentration of 5 mM for 30 minutes at room temperature (RT) with shaking (1000 rpm) on a Thermomixer (Thermo Fisher). Free cysteine residues were alkylated with 2-iodoacetamide at a final concentration of 10 mM for 30 minutes at RT in the dark. The reaction was quenched by adding DTT at a final concentration of 5 mM, followed by incubation at RT for 15 minutes. LysC was added at an enzyme-to-protein ratio of 1:200, and the mixture was incubated for 1 h at RT with shaking at 1000 rpm. Urea was then diluted to 2 M by adding 50 mM ammonium bicarbonate (ABC), and digestion with trypsin was performed at an enzyme-to-protein ratio of 1:100 at 37°C with shaking at 1150 rpm overnight.

After digestion, the peptide mixture was acidified to pH <3 by adding 50% trifluoroacetic acid (TFA), and desalted using 3-plug C18 stage tips (3M Empore^TM^ high-performance extraction disks). The stage tips were conditioned sequentially with i) 100 μL of methanol, ii) 100 μL of 70% acetonitrile (ACN)/0.1% TFA, iii) 100 μL of 0.1% TFA, iv) 100 μL of 0.1% TFA. After conditioning, the acidified peptide digest was loaded onto the stage tip. The stationary phase was washed with 100 μL of 0.1% formic acid (FA), and the peptides were eluted twice with 50 μL of 70% ACN/0.1% FA. Eluted peptides were dried under vacuum and reconstituted in 12 μL of 0.1% FA, followed by sonication and transfer to an autosampler vial. Peptide yield was quantified using a NanoDrop (Thermo Fisher).

### Mass Spectrometry parameters

Peptides were separated on a 25 cm, 75 μm diameter column with a 1.7 μm particle size of C18 stationary phase (IonOpticks Aurora 3, 1801220), using a gradient from 2% to 95% Buffer B over 90 minutes. Buffer A consisted of 0.1% FA in HPLC-grade water, and Buffer B consisted of 99.9% ACN, 0.1% FA. The separation was performed at a flow rate of 300 nL/min on a NanoElute 2 system (Bruker). MS data were acquired on a TimsTOF HT (Bruker) equipped with a Captive Spray ion source (Bruker) using a data-independent acquisition PASEF method (diaPASEF). The mass range was set from 100 to 1700 *m/z*, with an ion mobility range of 0.60 V.s/cm^2^ (collision energy: 20 eV) to 1.6 V.s/cm^2^ (collision energy: 59 eV), a ramp time of 100 ms, and an accumulation time of 100 ms. The dia-PASEF method covered a mass range of 400.0 to 1201.0 Da, a mobility range of 0.60-1.60 V.s/cm^2^, with an estimated cycle time of 1.80 seconds. The dia-PASEF windows were set with a mass width of 26.00 Da, a mass overlap of 1.00 Da, and 32 mass steps per cycle.

### Data Independent Acquisition (DIA) Data Analysis

Raw data files were processed using Spectronaut version 18.0 (Biognosys) and analyzed with the PULSAR search engine against a *Homo sapiens* UniProt protein database downloaded on 2023/05/25 (229,885 entries). Cysteine carbamidomethylation was specified as a fixed modification, while methionine oxidation, protein N-terminal acetylation, and deamidation (NQ) were set as variable modifications. A maximum of two trypsin missed cleavages was allowed. Searches used a reversed-sequence decoy strategy to control the peptide false discovery rate (FDR), with a threshold of 1% FDR for identification.

Inferred gene-level protein expression from LC-MS data was used for downstream analysis. For genes for which the inferred expression was discordant: (1) genes were excluded if the difference between the maximum and minimum absolute log_2_ fold change was >0.5, (2) for genes for which that difference was ≤0.5, the result yielding the largest absolute log_2_ fold change was used for downstream analysis. For the volcano plot, a log_2_ fold change threshold of >1 and q-value threshold of <0.05 were used. For gene set enrichment analysis (GSEA), the -log_10_(q-value)*sign(log_2_-ratio) was used as the ranking metric. The Hallmark gene sets^75^, ATF4 gene sets derived from the Molecular Signatures Database (MSigDB)^76^, and a manually curated gene set (“Key ATF4 targets”) reflecting ATF4 activity were used for GSEA. All the ATF4 gene sets used in this study are shown in Supplementary Table 7. An adjusted p-value of <0.05 was used as the threshold for significance for GSEA. All analyses were done using R 4.4.2. GSEA was run using the fgseaMultilevel function from the fgsea package^77^. The proteomic data are provided in Supplementary Table 6 and 7.

### RNA isolation and RT-qPCR

RPE mtDNA^ΔScal^ cells (5 × 10^4^) were seeded in 12-well plates in the complete medium, as described for cell maintenance. The following day, cells were washed with PBS, and the medium was replaced with experimental medium (DMEM supplemented with 10% dialyzed FBS, 100 units/mL penicillin, 100 µg/mL streptomycin, 1 mM pyruvate and 200 µM uridine) with additional 100 µM asparagine as indicated. After 24 hours, total RNA was extracted using an RNA purification kit (Norgen Biotek, 37500), according to the manufacturer’s protocol. cDNA was synthesized from total RNA using a cDNA Synthesis Kit (Bio-Rad, 1708891). Quantitative PCR (qPCR) was performed using Power SYBR^TM^ Green PCR Master Mix (Thermo Fisher Scientific, 4367659) with gene-specific primers (listed in Supplementary Table 4) on the QuantStudio^TM^ 7 Flex System. Target gene expression was normalized to *RPL19* mRNA levels.

### Tumor Xenograft assay

All animal procedures were performed in accordance with the MSKCC Institutional Animal Care and Use Committee (IACUC) guidelines. Subcutaneous injections of 2 × 10^6^ NCI-237 cells were administered into the flanks of 8-week-old male NOD.Cg-*Prkdc^scid^ Il2rg^tm1Wjl^*/SzJ mice (NSG^TM^, Jackson Laboratory). Two weeks post-injection, the mice were randomly divided into two groups. The control group received intraperitoneal (i.p.) injections of 150 µL PBS per mouse per day, while the asparaginase treatment group was injected with 80 IU asparaginase dissolved in 150 µL PBS per mouse per day. Tumor growth was monitored by caliper measurements every other day, beginning on the day of treatment. Tumor volume was calculated using the formula: length × width^2^ × 0.5. At the designated time points, the animals were euthanized via CO_2_ inhalation.

### Statistics and reproducibility

Unless otherwise specified, statistical analyses were performed using GraphPad Prism software. Data are presented as mean values, with error bars indicating the standard deviation and a minimum of three biological replicates (n ≥ 3), unless otherwise indicated in figure legends. The results shown are representative of at least two independent experiments. Statistical significance between two groups was determined using two-tailed Student’s t-test. One-way ANOVA was used for comparisons among three or more groups with a single independent variable. Two-way ANOVA was used for comparisons involving two independent variables (e.g., treatment and cell line). Additional details on statistical tests can be found in the figure legends.

### Figure Construction

Graphs were generated using GraphPad Prism (v.10) and compiled into final figures with Adobe Illustrator 2025. Chemical structure illustrations were produced with ChemDraw (v.22.0.0). Proteomic data shown in Figures 3a, 3b, and Extended Data Figure 3a were generated by R 4.4.2. Schematics shown in Figures 2i, 7e, 8f, and Extended Data Figures 1a were designed using BioRender.

## Data availability

Materials used in this study are available upon request. Further information and requests for resources and reagents should be directed to and will be fulfilled by the lead contact, Craig B. Thompson. The raw proteomic data files generated in this study have been deposited in PRIDE.

## Acknowledgements

We thank members of the Thompson laboratory, especially Tullia Lindsten and former members Natalia N. Pavlova (University of Utah) and Daphne Baker, for discussions and feedback. We acknowledge the use of the Integrated Genomics Operation Core, funded by the NCI Cancer Center Support Grant (CCSG, P30 CA008748), Cycle for Survival, and the Marie-Josée and Henry R. Kravis Center for Molecular Oncology. We acknowledge the help from the Proteomics Innovation Laboratory with proteomic analysis. We acknowledge the Weill Cornell Medicine Proteomics and Metabolomics Core Facility for performing LCMS analysis. We acknowledge the Cell Metabolism Core Facility at MSKCC. In addition, this work was supported by a grant from the NCI (R35 CA283988). Work in the A.S. lab is supported by a grant from NIH/NIA (R01AG085782). Z.B. reports support from the National Cancer Institute Cancer Center Core (grant P30-CA008748) supporting Memorial Sloan Kettering Cancer Center and from the National Cancer Institute’s Clinical Scholars Biomedical Research Training Program (T32CA009512-35) and The Alan and Sandra Gerry Metastasis and Tumor Ecosystems Center (GMTEC) Shulamit Katzman Endowed Postdoctoral Research Fellowship. R.C. is a Marie-Josée Kravis Women in Science Endeavor Graduate Student Fellow.

## Author information

### Contributions

R.C. and C.B.T. conceived the study. R.C. performed most experiments and analyzed the data. K.W.R. oversaw the design and execution of the experiments and assisted in the analysis of the data. Y.F. and T.K. provided key experimental materials under the supervision of A.S. Z.B. performed proteomic analysis. D.L. assisted in methods development. R.C., K.W.R., and C.B.T. interpreted the results and wrote the manuscript. All authors participated in discussing and finalizing the manuscript.

### Corresponding authors

Correspondence to Craig B. Thompson.

### Ethics declarations

C.B.T. is a founder of Agios Pharmaceuticals. He is on the Board of Directors of Regeneron and Charles River Laboratories. A.S. is a co-founder, consultant, and shareholder of Repare Therapeutics. The other authors declare that they have no competing interests. Z.B. reports honoraria from UpToDate (two chapters), the Fund for Innovation in Cancer Informatics (grant review), and the American Society for Clinical Oncology (associate editor for JCO CCI). Z.B. holds a provisional patent related to the prediction of oncologic systemic therapy toxicities. All COIs are unrelated to the current study.

